# Cooperative coding of continuous variables in networks with sparsity constraint

**DOI:** 10.1101/2024.05.13.593810

**Authors:** Paul Züge, Raoul-Martin Memmesheimer

## Abstract

A hallmark of biological and artificial neural networks is that neurons tile the range of continuous sensory inputs and intrinsic variables with overlapping responses. It is characteristic for the underlying recurrent connectivity in the cortex that neurons with similar tuning predominantly excite each other. The reason for such an architecture is not clear. Using an analytically tractable model, we show that it can naturally arise from a cooperative coding scheme. In this scheme neurons with similar responses specifically support each other by sharing their computations to obtain the desired population code. This sharing allows each neuron to effectively respond to a broad variety of inputs, while only receiving few feedforward and recurrent connections. Few strong, specific recurrent connections then replace many feedforward and less specific recurrent connections, such that the resulting connectivity optimizes the number of required synapses. This suggests that the number of required synapses may be a crucial constraining factor in biological neural networks. Synaptic savings increase with the dimensionality of the encoded variables. We find a trade-off between saving synapses and response speed. The response speed improves by orders of magnitude when utilizing the window of opportunity between excitatory and delayed inhibitory currents that arises if, as found in experiments, spike frequency adaptation is present or strong recurrent excitation is balanced by strong, shortly-lagged inhibition.

**Author summary:** Neurons represent continuous sensory or intrinsic variables in their joint activity, with rather broad and overlapping individual response profiles. In particular there are often many neurons with highly similar tuning. In the cortex, these neurons predominantly excite each other. We provide a new explanation for this type of recurrent excitation, showing that it can arise in a novel cooperative coding scheme that minimizes the number of required synapses. This suggests the number of required synapses as a crucial constraining factor in biological neural networks. In our cooperative coding scheme, neurons use few strong and specific excitatory connections to share their computations with those neurons that also need it. This way, neurons can generate a large part of their response by leveraging inputs from neurons with similar responses. This allows to replace many feedforward and less specific recurrent connections by few specific recurrent connections. We find a trade-off between saving synapses and response speed. Theoretical estimates and numerical simulations show that specific features of biological single neurons and neural networks can drastically increase the response speed, improving the trade-off.

## Introduction

The brain encodes continuous sensory or intrinsic variables in the coordinated activity of populations of neurons. The tuning curves (response profiles) of individual neurons in such populations are rather broad, leading to large overlaps between them [1, 2]. Further, there are often many neurons with highly similar tuning. Neuron populations with such features include simple cells in the primary visual cortex (V1) [3, 4], head direction cells in the anterior thalamic nucleus [5], tactile neurons in primary somatosensory cortex [6], place cells in the hippocampus [7] and grid cells in the medial entorhinal cortex [8, 9]. In machine learning, convolutional networks have overlapping receptive fields (RFs) that tile the input space [10]. RFs similar to those in visual cortex emerge by learning a sparse code for natural images [11], and RFs similar to grid cells emerge through training on navigation tasks [12, 13].

Neurobiological data show that neurons with strongly overlapping receptive fields are predominantly excitatorily coupled: Synaptic connections between similarly-tuned excitatory principal neurons are more likely [14], stronger and more often bidirectional [15, 16]. In line with this, the strongest incoming synapses provide excitation that matches a neuron’s RF [16, 17]. Furthermore, highly similarly tuned principal neurons have overall, i.e. including indirect, polysynaptic connections, a net excitatory effect on each other [18, 19]. In contrast, if the tuning is barely similar or dissimilar, the net effect is inhibitory.

Such recurrent excitatory connectivity may seem unintuitive from a normative standpoint, as it amplifies noise [20] and can increase response times [21, 22]. Previous studies suggested that it may support persistent activity and thus working memory [23, 24] and that it may implement complicated priors [25].

Neural networks, however, evolved subject to physiological and physical constraints [26–29], including metabolic cost and available space. Optimizing for specific features can largely determine the neural network and lead to solutions that are in other aspects sub-optimal. A prominent example for this is a recent version of the efficient coding hypothesis [30–34]. It posits that neural networks greedily minimize the number of used spikes or the rate activity, which contribute to metabolic cost. The network connectivity obtained from the optimization is, however, very dense, which is not found in experiments. Further, the coding scheme is “competitive”, in the sense that similarly tuned neurons compete for the opportunity to generate spikes. In other words, such neurons take away spikes and activity from each other. This predicts inhibitory couplings between very similarly tuned neurons, contrary to the experimentally observed physiological and effective excitatory interconnectivity between them.

Here, we explore the implications of “cooperative coding” in a neural network. In this newly proposed scheme, neurons avoid replicating computations through feedforward weights whose results are already accessible from the activity of other feature neurons. Instead, each feature neuron performs only a non-redundant feedforward computation. It then achieves the required response by additionally incorporating the results already obtained by similarly tuned feature neurons through recurrent connections. In other words, feature neurons do not independently replicate shared parts of the computations through feedforward weights, but they transmit them through recurrent connections to each other. The resulting connectivity is like-to-like, i.e. strong and effectively excitatory between similarly tuned principle neurons, as observed in experiments. Interestingly the scheme optimizes the number of synapses in a network, while maintaining the required neural network dynamics. Such an optimization differs from the common focus on saving spikes and may be imposed by space restrictions or cost of maintaining synapses [28, 35].

## Results

To demonstrate the concept of cooperative coding, we consider a layer of feature neurons (output neurons), which receive feedforward input from an input layer as well as recurrent input. The task of the feature neurons is to generate a weighted sum of the inputs with weight strengths that decay exponentially with the distance of an input from the preferred input. We assume that the functionally relevant network response, representing the desired features (outputs), is the steady state activity. The desired outputs are linear functions of the inputs. Neural responses can hence be characterized by linear RFs and implemented by feedforward connectivity alone. Importantly, they can also be implemented using mixtures of feedforward and recurrent input.

We will compare the different network implementations in terms of the space requirement, approximated by the number of required synapses, and in terms of the metabolic cost to keep up the stationary state. Finally we will compare the response times and demonstrate how they can be substantially decreased in networks with spike frequency adaptation (SFA) or balancing inhibition.

### Models

In the following, we analyze three concrete examples of cooperative coding: (i) encoding a one-dimensional stimulus, (ii) simultaneously encoding two one-dimensional stimuli with linear mixed selectivity (MS) and (iii) encoding a two-dimensional stimulus. For ease of description, we focus on translationally invariant RFs. (Approximate) translational invariance, meaning that offset RFs have similar shapes, is a common characteristic of experimentally encountered RFs [2, 4, 5, 8, 9]. Further, it is a common characteristic of RFs that emerge in machine learning [10, 11]. Although the RFs that we consider do not have the precise shape of measured RFs, for example those of simple cells in V1 [36], they share the key properties of localized, overlapping and broadening RFs that tile the represented space. Indeed, RFs of neurons in hierarchically higher layers are often broader and constructed from those in lower layers [37, 38].

### Encoding a 1D stimulus

As a concrete, analytically tractable model that illustrates how cooperative coding works and can save synapses, we consider RFs that tile the one-dimensional parameter space of a stimulus (see Fig. 1a)). An input neuron *j, j* = 1, …, *N* signals the presence and strength of a stimulus with a specific parameter *j* by nonzero activity *r*_*j*_ *>* 0. The task of the feature layer is to generate a response that is maximal at the preferred stimulus parameter and then decays exponentially the more different the stimulus becomes from the preferred one. This behavior is qualitatively similar to commonly observed tuning curves such as orientation tuning curves or place fields. We further assume that if multiple stimuli are present, the feature layer responses to their different parameters superpose linearly.

**Fig 1:**
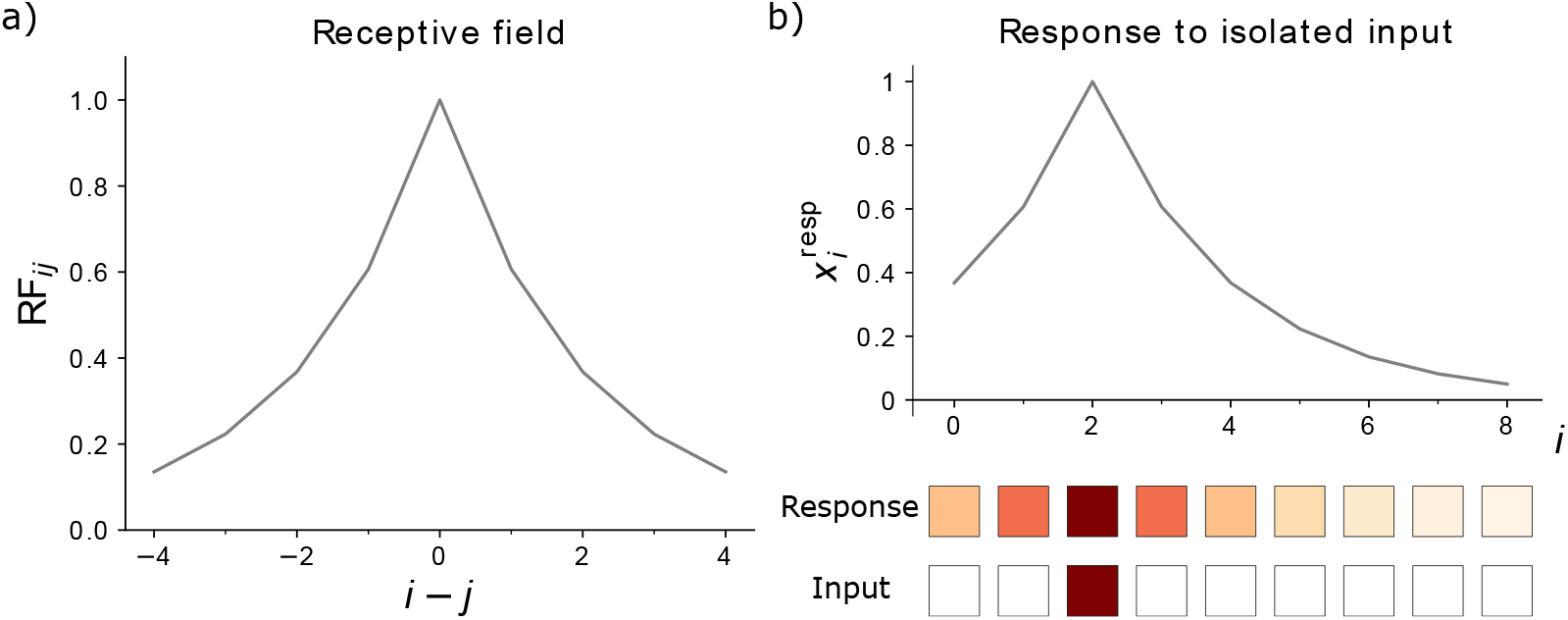
Receptive field and response of 1D network. **a)** The RF of neuron *i* is peaked at *j* = *i* and decays exponentially with |*i* −*j*|. The receptive field width parameter is *d* = 2. **b)** Top: The network response to an isolated unit input, here located at *j*_0_ = 2, has the same shape and amplitude as a neuron’s RF. It peaks at *i* = *j*_0_. Bottom: Input and feature neurons are shown with color-coded activities *r*_*j*_ and *x*_*i*_, respectively. Increasingly dark red color represents higher activity; white squares indicate inactive neurons.

As an example, the inputs may be interpreted as a simple model for the representation of the orientation of a bar in the early visual system. *r*_*j*_ *>* 0 then means that the orientation is within the *j*th bin of the total orientation range [0, 180^°^]. The transformation from input to features in our model describes the combination of responses from hierarchically lower visual areas to hierarchically higher ones [39]. As another example, the neurons may model the activity of place cells on a periodic, closed track. The transformation then models the transformation from input neurons with smaller place fields to neurons with larger place fields. Such a transformation may take place from the hippocampal dentate gyrus to the downstream area CA3 [40]. In our model, the input generates a simple encoding of the current location, where input neuron *j* is active if the animal is in the *j*th location.

The desired stationary feature layer activity can be expressed as

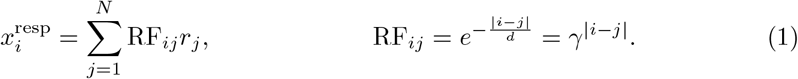

Here *r*_*j*_ is the activity of the *j*th input neuron, *γ* = exp(−1*/d*) and *d* defines the width of the RF. We use periodic boundary conditions. For computations with neuron indices, this means that |*i* − *j*| means min_*n*∈{−1,0,1}_ |*i* − *j* + *nN* |. There are as many input as feature neurons. We note that, because of the symmetry RF_*ij*_ = RF_*ji*_, the vector RF_*k*·_, describing the RF of feature neuron *k*, is the same as RF_·*k*_, the network response when only input neuron *k* is active, compare Fig. 1a) and b). Summarized in a formula, we have 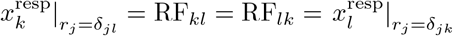, where *k* is fixed and *l* variable.

### Feedforward implementation

To model the temporal dynamics of the neurons, we choose a standard simple linear rate network model [41, 42]. The purely feedforward network that generates the response eq. (1) as stationary state is then given by

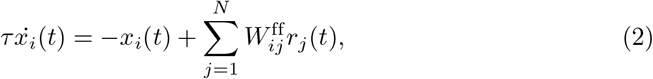

with 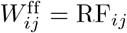 and a time constant *τ*. In the stationary state, we have 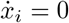 for all *i* and thus

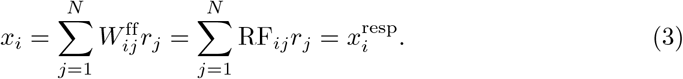

This state is asymptotically stable and globally attracting; the flow is a contraction to it. These properties follow immediately from the fact that the system is linear and has a unique fixed point, which is asymptotically stable because all eigenvalues of the matrix specifying the homogeneous differential equation are negative, equal to − 1*/τ* [43, 44]. *To approximate the network with a characteristic number of feedforward weights that is smaller than N*, we require synapses 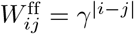 only where |*i* − *j*| ≤ *d*. This defines the RF size *n*_RF_ = 2*d* + 1 as the number of feedforward synapses per neuron needed to implement the RF within a distance *d* around its center.

### Cooperative implementation

The same stationary neuronal responses can be obtained as the steady state of a recurrent network that uses cooperative coding. It requires only three synapses per feature neuron, two recurrent and one feedforward one. This network’s dynamics are given by

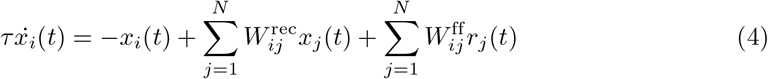

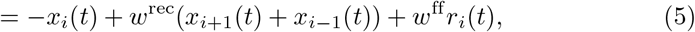

with weights 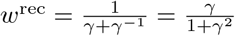 and 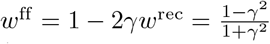. If the receptive fields are not narrow (*d* is not small against 1), the two recurrent connections are strong, in the sense that *w*^rec^ is not small against 1. Thus the network features strong like-to-like excitation and is driven by feedforward input. One can straightforwardly verify that 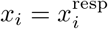 is indeed a stationary state of the network, by inserting eq. (1) into eq. (5), see appendix S1. The reason for this is ultimately that the desired response of a neuron *i* can be largely generated by summing the responses of the two neurons *i* ± 1 neighboring *i*, see eq. (9) and Fig. 2b). This is achieved by the recurrent connections. The missing part is contributed by the feedforward input. This state is asymptotically stable as all real parts of the eigenvalues of the matrix defining the homogeneous system are negative, see appendix S1. For broad receptive fields (where *γ* ≲ 1), the recurrent connections are nearly as strong as possible: their sum 2*w*^rec^ is close to 1, the value beyond which the network becomes unstable. The stationary state is also the only stationary state. Since the system is linear, the state is therefore a global attractor as for the feedforward network [43, 44]. Thus, for constant input the network forms this stable response pattern.

**Fig 2:**
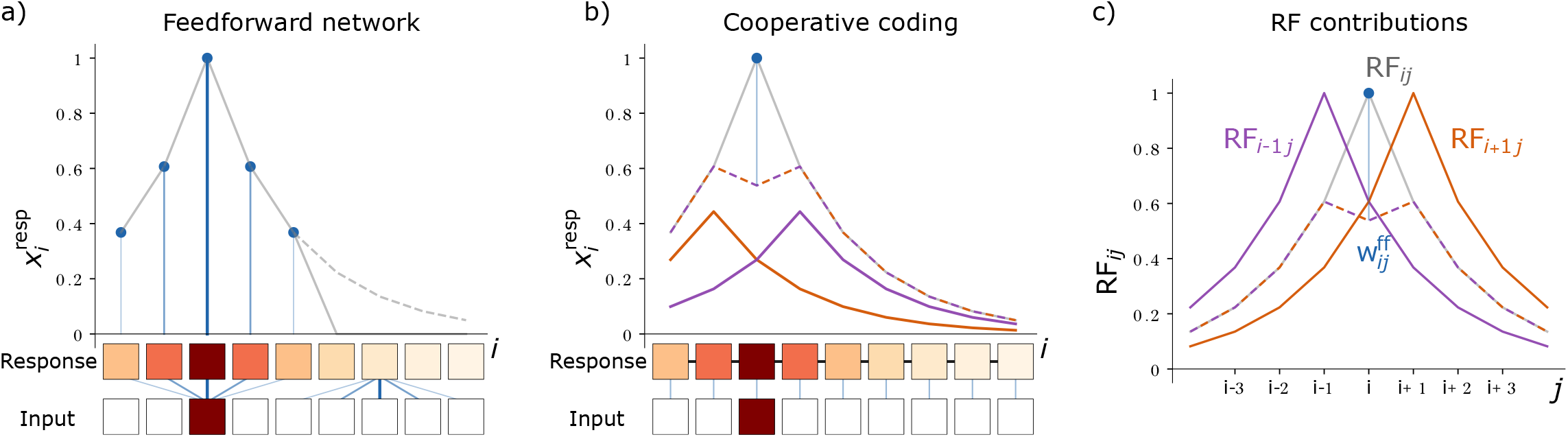
Schematics of feedforward and cooperatively coding networks. **a)** Top: In the feedfoward network, the response 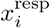 (gray solid curve) to an isolated input is fully generated by the neurons’ feedforward inputs (blue lines and dots). For the displayed RF width *d* = 2, five neurons receive feedforward input, so that the network response (gray solid curve) represents ≈ 63% of the summed target response (gray dashed curve). Bottom: Feature and input neuron activities as in Fig. 1. Outgoing feedforward synapses from the active input neuron *j* = 2 and incoming feedforward synapses to feature neuron *i* = 6 are shown in blue. **b)** Top: In the cooperatively coding network model, the network response (gray solid curve) is the sum of feedforward input (blue line and dot) and recurrent input (brown-purple dashed curve). For the displayed case of an isolated input, only one neuron receives feedforward input, which induces a part of the stationary response of the most active feature neuron. The rest of the response and all other responses are induced by recurrent input from neighboring neurons. The total recurrent input that each feature neuron receives is the sum of recurrent input from the right (brown solid curve) and left neighbor (purple solid curve). Bottom: Each feature neuron receives one feedforward synapse (blue lines) and two recurrent synapses (black lines, all recurrent connections are bidirectional). **c)** The RF of feature neuron *i* (RF_*ij*_ for varying *j*, gray solid curve) is the weighted sum (brown-purple dashed curve) of the RFs of its left (RF_*i*−1 *j*_, purple) and right neighbors (RF_*i*+1 *j*_, brown) plus a contribution from feedforward input (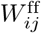, blue line and dot). All shown RFs have width *d* = 2.

### Cooperative coding

Cooperative coding can be understood as sharing of the information that an individual neuron obtains from external input specifically with those neurons that also need it. This allows to generate most of the neuronal responses from sparse recurrent connectivity. Especially very similarly tuned neurons will project strongly excitatorily onto each other; oppositely tuned neurons would inhibit each other.

As a concrete example, we introduced the networks eq. (5), where it suffices that each neuron receives input from only one input and two feature neurons. Still, each neuron effectively responds to 𝒪 (*d*) input neurons. This is possible because the feature neurons recurrently share their activity, and hence their access to feedforward input, with their neighbors. These in turn share it with their neighbors, thus propagating it through the network. The network response then forms dynamically through the interplay of feedforward input and recurrent interactions.

The coding relies on the fact that despite very few feedforward and recurrent synapses, poly-synaptic connectivity can still be far-reaching [45, 46]: For further clarification of this point consider the approximate, discretized dynamics [47, 48]

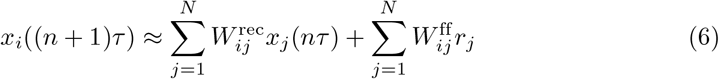

that lead to the same steady state as the time-continuous dynamics eq. (4). The response to a constant input *r* after *n* time constants is

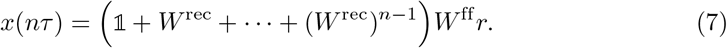

It is determined by *W* ^rec^ and its higher powers, which reflect the redistribution of feedforward input via poly-synaptic recurrent pathways.

The coding scheme can also be understood as feedforward inputs providing a correction to the response that is mainly constructed from the sparse recurrent input. To clarify this we focus on elementary stationary responses, namely those that are driven by a single unit input from neuron *j*; the input activity is *r*_*k*_ = *δ*_*kj*_. Responses to more complicated input patterns are weighted linear sums of such elementary responses. Consider feature neuron *i* and assume that all other neurons already respond correctly. The stationary activity of neuron *i* in response to a single unit input from neuron *j* is then RF_*ij*_, while those of the other network neurons *k* is RF_*kj*_. Equation (4) with 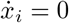 implies that RF_*ij*_ is the sum of the RFs of its presynaptic feature neurons and its feedforward connectivity,

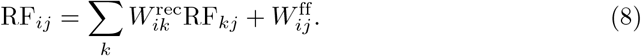

For the specific network eq. (5) we have

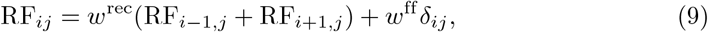

illustrated in Fig. 2c). In this network, the weighted and summed responses of neuron *i*’s nearest neighbors are thus already very close to neuron *i*’s target response. This is enabled by the specific exponential shape of the RFs. Only for the preferred input of a feature neuron, the response is too low. The neuron corrects for this by recruiting the missing input through a feedforward connection. Such an “explanatory gap” that is left by the recurrent inputs and can be filled by external input is important for cooperative coding, because the output must depend on external input.

### Spatial demand and metabolic cost

To compare the efficiency of the introduced implementations, we focus on two cost dimensions: the space needed to implement the network and the metabolic cost of generating the stationary dynamics. As measure for the space needed for the network we take the number of synaptic connections, or, in other words, the L0 norm of the synaptic weight matrix. In the feedforward network eq. (2), it increases linearly with the width *d* of the RF (if small responses can be neglected). In the recurrent network eq. (5) three synapses per neuron suffice to generate the desired stationary response regardless of the RF size. We show in appendix S1 that the recurrent network eq. (5) therefore minimizes the L0 norm.

For the metabolic cost of generating the stationary dynamics, we can take a multiple of the L1 norm of the synaptic currents (see Discussion). Since in both networks all modeled synaptic currents are excitatory, the L1 norm of synaptic currents equals the total synaptic current. In the stationary state this current is the same in both implementations, because neurons have the same stationary activity; therefore also the cost is the same. We conclude that the metabolic cost for maintaining the stationary network state is the same in both the feedforward and the recurrent network implementation.

### Response speed

In the feedforward network, activity converges with the intrinsic time constant *τ*, which we define as its response time. In the cooperatively coding network, the excitatory recurrent connectivity increases the response time: Fig. 3 shows the dynamics of the response formation. The L1-norm of the deviation of the response from the steady state,

**Fig 3:**
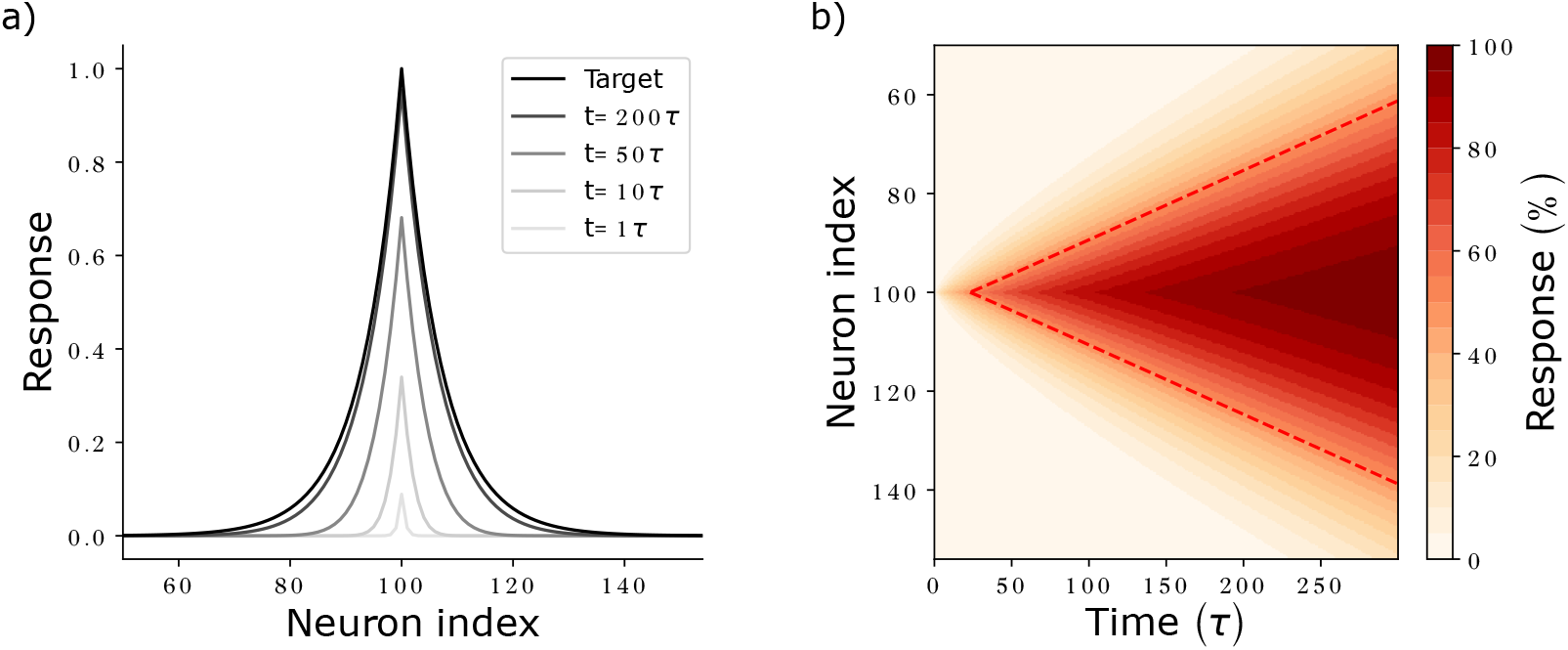
Response formation and activity propagation. **a)** Network activity at different times (shaded curves) after *r*_100_ has been set from 0 to 1. For long times, network activity approaches the target response (black curve). **b)** Development of the activity of neurons (y axis) with time (x axis), measured relative to their target activities. The diagonal fronts of equal relative activities indicate propagation of activity with constant propagation speed. The time points at which neurons reach 50% of their final activity are connected by a red dashed line. Parameters: *w*^rec^ = 0.5 · 1*/*(1 − 1*/*100), such that *τ*_resp_ = 100*τ* (see eq. (11)), *N* = 200 neurons.

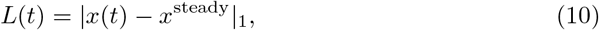

which we use as a loss measure, decays exponentially with time constant

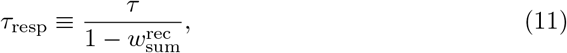

(see Fig. 4), where 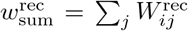 is the sum of recurrent weights arriving at (or, equivalently, originating from) a neuron. We define *τ*_resp_ as the response time of the networks. It scales inversely with the difference of the largest eigenvalue of the network, which is equal to 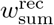 (cf. appendix S1), from 1. In particular, it depends only on the summed recurrent weights. Equation (11) holds generally, for networks of the type eq. (4) with purely excitatory circulant recurrent weight matrix and convergent dynamics.

**Fig 4:**
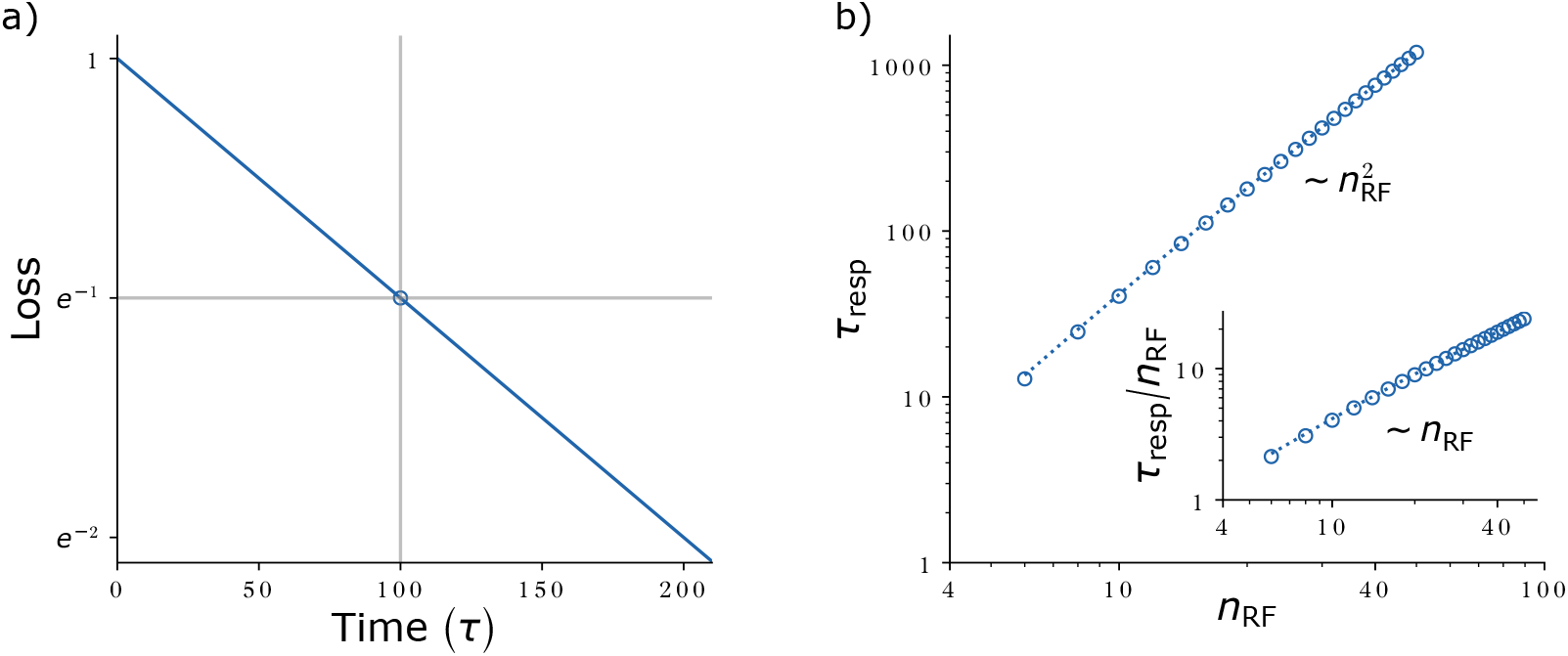
Loss evolution and response speed - synapse number trade-off. **a)** Exemplary loss evolution for a network with 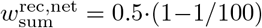 so that *τ*_resp_ = 100*τ*. Experimentally, *τ*_resp_ is determined as the time (gray vertical line) at which the loss drops to e^−1^ (gray horizontal line, blue open circle). **b)** Response times *τ*_resp_ (circles: simulation results; dotted line: analytical solution eq. (12)) for target RFs of different widths *n*_RF_. Data was created by scanning *n*_RF_, setting 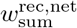 to yield a RF of size *n*_RF_ and determining *τ*_resp_ from the loss dynamics.

We now specialize the result to networks with nearest-neighbor coupling eq. (5) that generate the RFs eq. (1). Inserting *w*^rec^ = 1*/*(*γ* + *γ*^−1^) and *γ* = exp(−1*/d*) relates the response time to the RF width. By approximating exp(±1*/d*) ≈ 1 ± 1*/d* + (1*/*2)*d*^2^ for large *d*, we obtain *w*^rec^ ≈ 1*/*(2 + 1*/d*^2^) and, inserting this into eq. (11),

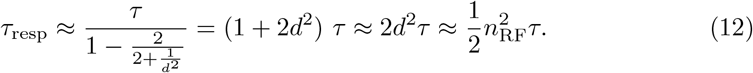

In the last part of the equation we used that *n*_RF_ = 1 +2*d* ≈ 2*d* for large *d*. Equation (12) shows that wide RFs require long equilibration time. This is because they need strong recurrent weights with a largest eigenvalue close to 1. Further the equation reveals the trade-off between response time and number of employed synapses: The feedforward implementation eq. (2) needs *n*_RF_ synapses and has a response time *τ*. The recurrent implementation thus saves *n*_RF_ − 3 ≈ *n*_RF_ synapses per feature neuron. Equation (12) shows that the response time increases quadratically in the number of saved synapses, see also Fig. 4b).

The quadratic dependence of the response time on *d* reflects that, as the RF becomes wider, not only does activity have to spread further, it also spreads more slowly: this is consistent with the idea that the settling of a neuron depends (indirectly) more on activity propagating back from more distant neurons that settle after it.

### Faster response with spike frequency adaptation

For activity to rapidly spread through the network, neurons need to be able to cause a large activity change in their neighbors within a short period of time. To achieve this, they need strong recurrent weights. However, recurrent weights are restricted to 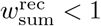 to not cause runaway activity. We now show how spike frequency adaptation (SFA) can help ease this conflict and speed up network dynamics. SFA is typical for excitatory principal neurons and induces a reduction of their response to constant inputs in the long run [42, 49, 50].

We model SFA through a negative-feedback adaptation current *u*(*t*), which is triggered by neuronal activity *x*(*t*) and characterized by its scale *a*_SFA_ and time constant *τ*_SFA_,

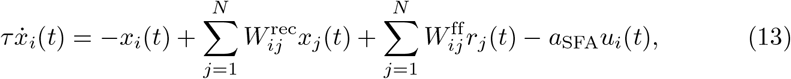

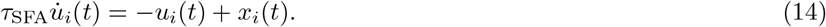

[51–53]. Setting 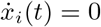 and 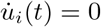 yields the steady state. We see immediately that it implies *u*_*i*_ = *x*_*i*_. Inserting this into eq. (13) shows that in the stationary state the spike frequency adaptation results in a stronger leak current, −(1 + *a*_SFA_)*x*_*i*_. Dividing by 1 + *a*_SFA_ yields

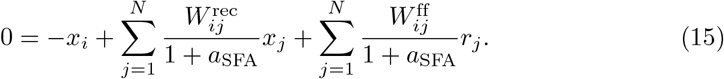

Consequently, in order to implement the same response as a network without SFA (*a*_SFA_ = 0, cf. eq. (4)), the recurrent and feedforward weights have to be scaled up by a factor of 1 + *a*_SFA_. The additional excitatory synaptic input compensates in the steady state the added inhibitory adaptation current.

To understand the network dynamics, it is instructive to consider the limit *τ*_SFA_ → 0 where *u*_*i*_(*t*) → *x*_*i*_(*t*) as in the steady state. Inserting this into eq. (13) and again dividing by 1+*a*_SFA_ yields an equation equivalent to eq. (4) with shortened neuronal time constant and increased weights,

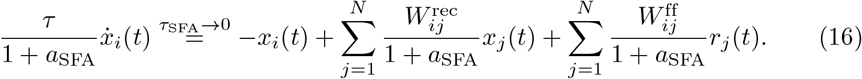

we see that a network with arbitrarily fast SFA and appropriately scaled weights has the same dynamics as a network without SFA, but with its time constant reduced by 1 + *a*_SFA_. This factor only depends on *a*_SFA_ and is independent of the RF width that the network implements. We might thus expect that introducing SFA with a given *a*_SFA_ and small *τ*_SFA_ causes a constant speedup, but still results in a quadratic dependence of 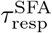 on *n*_RF_ (see eq. (12)).

There is, however, an additional possibility: SFA might yield faster dynamics for finite, nonzero *τ*_SFA_. This is because then *u*_*i*_(*t*) lags behind *x*_*i*_(*t*), which creates a temporal ‘window of opportunity’. Within this window, the up-scaled weights can mediate strong interactions that are not yet cancelled by the retarded adaptation currents of the receiving neurons. In our networks, this leads to the following concept to exploit SFA: During the initial response phase, strong weights should cause a fast response while SFA keeps the steady state before and after an input change at the desired activity values as well as dynamically stable. In particular, the modified recurrent synaptic weights may then be (and to optimally exploit SFA: should be) so strong that without the SFA current the network dynamics are unstable.

This second possibility applies to our networks: Measuring the response time as a function of *τ*_SFA_, we observe that it first decreases when increasing *τ*_SFA_ from zero and reaches a minimum at a nonzero, optimal value of the SFA time scale (appendix S4). Increasing *τ*_SFA_ further eventually causes diverging activity, because the retarded adaptation current *u*(*t*) becomes so slow that it never compensates the stronger input due to the up-scaled weights. We note that also when keeping *τ*_SFA_ at a fixed value, there is an optimal nonzero value of the inhibitory feedback strength *a*_SFA_. Importantly, we find that introducing SFA with finite *τ*_SFA_ and optimal *a*_SFA_ improves the scaling of 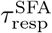 with *n*_RF_ from quadratic as without or with arbitrarily fast SFA to linear.

To incorporate SFA in a cooperatively coding network, we modify the weights in eq. (5) as described above and add the SFA current. For the neuron activities, this yields the dynamical equation

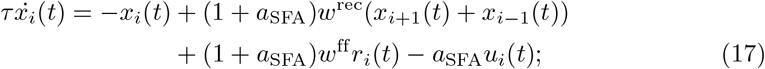

where the adaptation current obeys eq. (14).

Concerning the use of resources, SFA does not require additional synaptic connections, so the spatial demand of the cooperatively coding network is the same as in the original model eq. (5). The increased weights, however, lead to stronger synaptic currents. Together with the added adaptation currents, this increases the energetic cost of maintaining the stationary state.

## Balanced networks

The networks we studied so far had only excitatory synapses, while biological neural networks also have recurrent inhibition, which balances the excitation [32, 54]. These are likely required for a range of reasons, such as ensuring network stability and maintaining irregular spiking activity [55–59]. Given their existence, we here show how inhibition can be used to speed up the network response, in an architecture that still relies on few synapses.

The individual excitatory and inhibitory currents can be much larger than their sum and precisely temporally balanced with a lag smaller than the neuronal time constant [60, 61]. Further, many inhibitory neurons are rather sharply tuned [61–63], sometimes similarly sharply as excitatory ones.

To incorporate inhibition consistent with these experimental findings, we add inhibitory interneurons to the existing network of excitatory feature neurons, which represent principle neurons. The interneurons derive their tuning from the feature neurons via specific recurrent inputs. For concreteness we assume that there are as many interneurons as feature neurons and that each interneuron follows the activity of one feature neuron with a small delay, *τ*_lag_. Equation (4) thus becomes

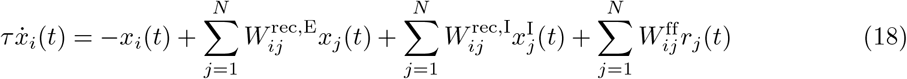

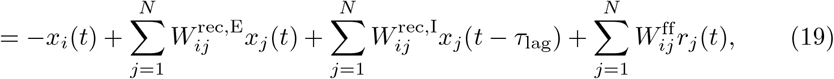

where 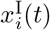 is the inhibitory activity, which equals the delayed excitatory feature neuron activity 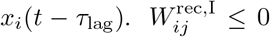 is the coupling from inhibitory neuron *j* to feature neuron *i*. We now introduce the state change of feature neuron *i* between *t* − *τ*_lag_ and *t*,

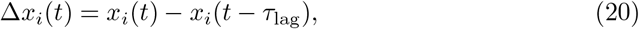

as well as the the sum of recurrent excitation and inhibition

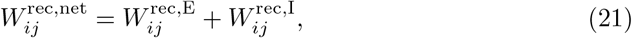

which we call net weights. These definitions allow to rewrite eq. (19) as

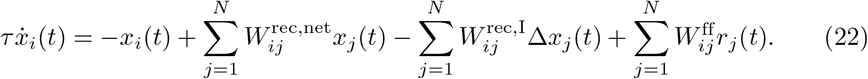

we now insert the values of the cooperatively coding network eq. (5) into the equation and further assume that the interneurons inhibit and balance the same sets of neurons that their driving feature neurons excited, i.e. 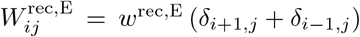 and 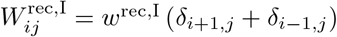. Equation (19) then becomes

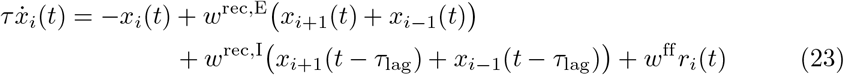

and eq. (22)

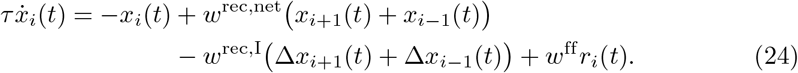

the residual inhibitory interaction, i.e. the inhibitory interaction that is not included in *w*^rec,net^, depends on Δ*x*_*i*±1_(*t*) and therefore only acts when there are activity changes during the preceding brief E-I lag. A total change *δx*_*j*_ in the activity of neuron *j* causes an integrated postsynaptic activity change in neuron *i* of 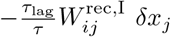 (see appendix S5).

To connect eq. (24) to our previous, unbalanced network eq. (5), we set

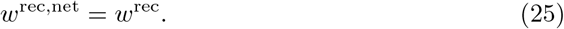

the dynamical equations then agree if *w*^rec,I^ = 0 or Δ*x*_*i*_(*t*) = 0. The latter is satisfied in the steady state. The steady state is thus independent of the amount of inhibition (given that excitation covaries with inhibition such that *w*^rec,E^ + *w*^rec,I^ = *w*^rec^ as implied by eq. (25)) and it is the same as in eq. (5); its stability can, however, change. We may thus think of *w*^rec,net^ as defining the RF, and of *w*^rec,I^ as affecting the dynamics. During the build-up of the response strong excitation ramps up slightly before the balancing inhibition. The analytical solution of eq. (24) and our stability analysis (see next section and Fig. 9) show that throughout this window of opportunity excitation may be up to approximately *τ/τ*_lag_ times larger than the net interaction without destabilizing the network. This strong interaction allows a much quicker propagation of activity, convergence to the steady state and decay of the loss function.

### Spatial demand and metabolic cost in the balanced network

Compared to the unbalanced network, the balanced network requires three additional synapses per principle feature neuron, one E-to-I and two I-to-E synapses, i.e. a total of six synapses. This is again independent of the RF width, such that for large RFs, the balanced, cooperatively coding network still saves synapses compared to the feedforward network. It requires additional space for the inhibitory neurons, which may, however, be needed for other purposes anyways.

Also the metabolic maintenance cost increases, since there are more neurons. Further, there is an increased metabolic cost to sustain the synaptic currents in the stationary state: In this state, large parts of the excitatory and inhibitory currents cancel to give rise to a net current that equals the one in the purely excitatory recurrent network, see eq. (21) and eq. (25). In the L1 norm of the synaptic currents, the excitatory currents and the absolute inhibitory currents, however, add up. The metabolic cost thus increases by the amount of excitatory and inhibitory currents that cancel each other.

### Response speed in the balanced network

Under some additional assumptions, the evolution of the L1-loss eq. (10) can be analytically approximated as the solution of a linear delay differential equation, see appendix S6 for details. The resulting dynamics are those of a damped oscillator, see Fig. 5a): For weak inhibition, they are ‘overdamped’ in the sense that they are well described by the sum of two exponentials with different decay rates. At a specific intermediate inhibitory strength, the two decay rates agree and we have ‘critical damping’. For stronger inhibition the dynamics are ‘underdamped’ in the sense that the loss behaves as the absolute value of an oscillation with exponentially decaying amplitude. Overly strong inhibition causes divergence of the dynamics.

**Fig 5:**
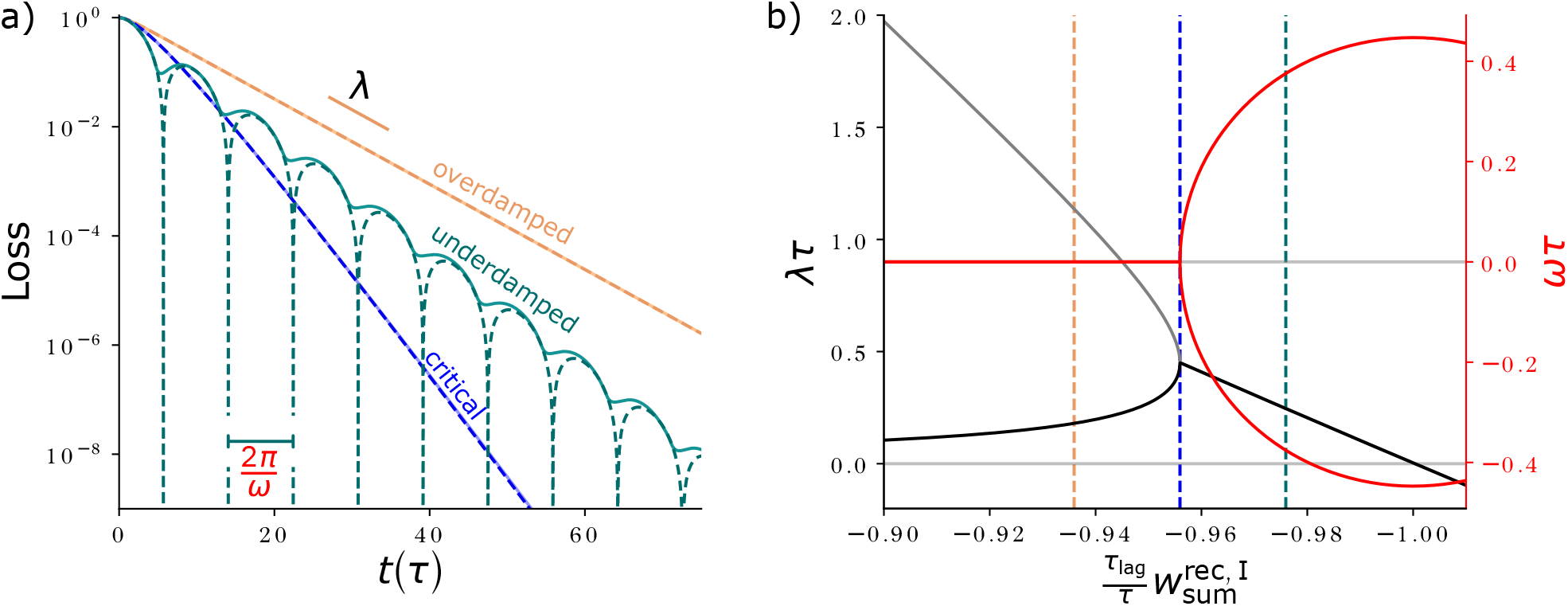
Loss evolution for different inhibitory strengths. **a)** Loss evolution (dashed: analytical approximation (cf. appendix S6, eqs. (64) and (74), partly occluded; solid: network simulation) for inhibitory strengths that are slightly weaker (orange), equal (blue) or slightly stronger (teal) than the critical strength, on a logarithmic scale. The slope of the decay is given by *λ* (see b), explicitly highlighted for the overdamped dynamics. The oscillation period of the underdamped dynamics is *T*_osci_ = 2*π/ω*. In case of oscillations, the analytic approximation briefly reaches zero loss once in a period (sharp dips in dashed curve). In the network simulation there is also a pronounced oscillation, but there always remains a finite error (solid, see appendix S6). **b)** Real part (decay rate *λ*, black/gray) and imaginary part (oscillation frequency *ω* times ± 1, red) of the complex frequency of the exponential loss evolution, scaled by *τ*. For weak inhibition there are two exponentially decaying modes (*λ*, black and gray curve). At the critical inhibitory strength (blue dashed vertical line) there is only a single decay rate and no oscillation. The decay rate (in the overdamped case: of the relevant slower-decaying mode) is maximized. For stronger inhibition, network activity begins to oscillate (nonzero *ω*, red), and diverges for 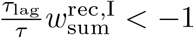, where *λ* becomes negative. Dashed vertical lines show the inhibitory strengths scaled by *τ*_lag_*/τ* for the curves in a) 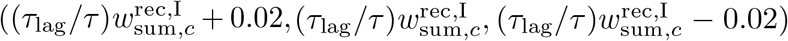. Parameters: 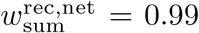, *τ* = 1, *τ*_lag_ = 0.1, and *N* = 200 for the network simulation.

In the overdamped regime, the smaller decay rate is the relevant one, as it dominates the speed of the decay for longer times. The larger decay rate rather describes how quickly faster dynamics, that may be present due to the initial conditions, are suppressed and the dynamics converge to the slower mode. The smaller decay rate increases when the strength of inhibition approaches its critical value. The same holds for the single decay rate in the oscillatory regime. At the critical inhibitory strength the overall decay of the loss is thus fastest, see Fig. 5b). We find analytically that its decay time constant is approximately proportional to the geometric mean of the response time in absence of inhibition and the inhibitory delay,

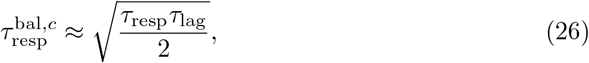

(cf. eq. (88) in appendix S6); the superscript “*c*” indicates that the result holds for critical inhibitory strength.

Importantly, this implies that the scaling of the response time with the receptive field width and size improves compared to the purely excitatory network. This is because 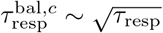. Inserting eq. (12) into eq. (26) for the 1D network, we obtain

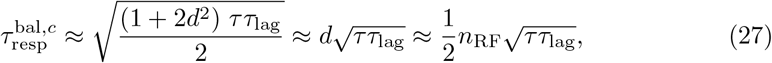

which is only linear in the RF width *d* and size *n*_RF_, instead of quadratic as in the case of 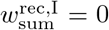, compare eq. (27) with eq. (12) and in Fig. 6a) the red and blue dotted curves. As a consequence also the speedup gained through the inhibition, 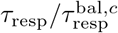, increases for wider RFs. We finally note that the interactions mediated by *W* ^rec,I^ can also be thought of as implementing an excitatory transmission of activity changes: an activity change in neuron *j* adds an activity change with the same sign to neuron *i*, because −*W* ^rec,I^ in eq. (22) is positive. Thus activity changes in different neurons in the network amplify each other. In the limit of small *τ*_lag_, the temporal derivative is transmitted, see appendix S5.

**Fig 6:**
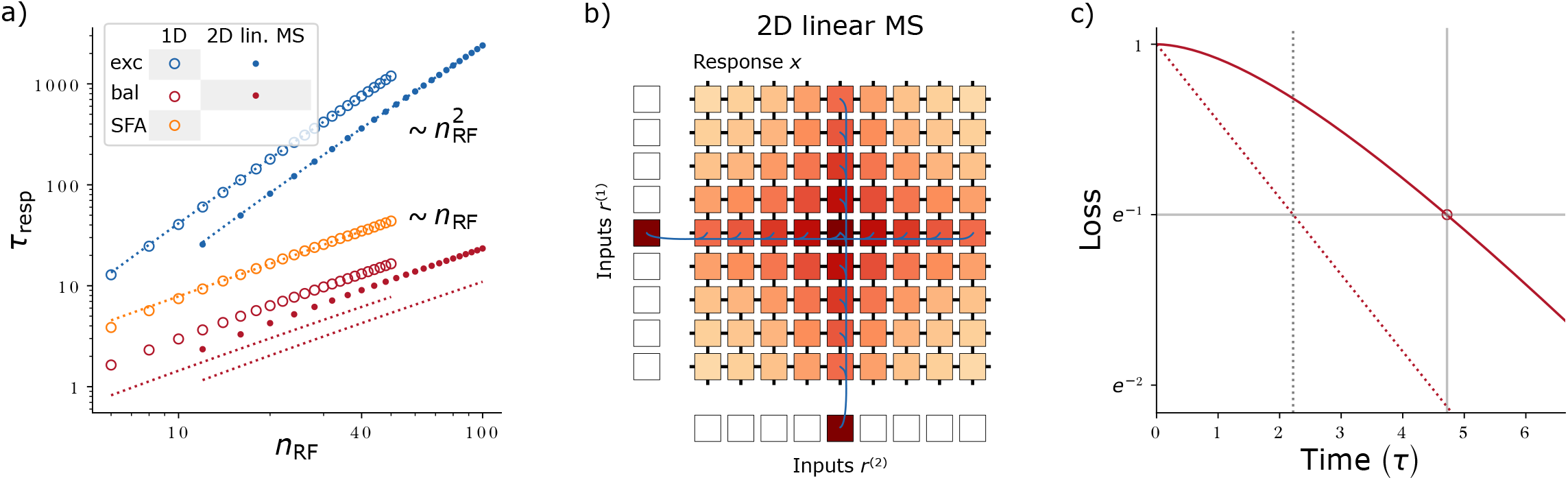
Response speed of networks with inhibition and linear MS. **a)** Response times for the 1D network and for the 2D linear MS network. The quadratic scaling of *τ*_resp_ with *n*_RF_ for the excitatory networks (blue) can be improved to a linear dependence by introducing balancing, delayed inhibition (red) or SFA (orange). Open (1D network) and filled (MS network) circles display numerical results. Alike colored dotted (1D network) or continuous (MS network) curves show theoretical estimates (eqs. (12), (27), (33) and (34)) or, for the SFA network, fit results (monomial fit: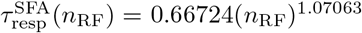). We use the slowest decaying eigenmode to theoretically estimate the response times (see c). Since the balanced networks are not initialized in this eigenmode (in contrast to the purely excitatory networks), the numerically measured response times (red markers) lie above the theoretical values (red lines). **b)** Schematic of a 2D network with linear MS. Feature neurons are arranged on a two-dimensional grid (labeled ‘Response *x*’). Each receives feedforward input from two arrays of input neurons (labeled ‘Inputs *r*^(1|2)^’) and four recurrent inputs. Feedforward and recurrent synapses are shown in blue (exemplarily) and black, respectively. Input and feature neuron activities are color-coded. The (linear) network response is the sum of the responses to input one and input two. **c)** Exemplary loss evolution for a 1D network with lagged inhibition. Due to the temporally constant initialization (*x*_*i*_(0) = 0, Δ*x*_*i*_(0) = 0), the network activity (solid red curve) converges initially more slowly than the network’s slowest eigenmode (dotted red line). The experimentally measured response time (continuous vertical gray line) is defined as the time when the loss has decayed by 1*/e* (red open circle, horizontal gray line, see also Fig. 4a). It is larger than that of the network’s eigenmode (dotted gray line), which we use as analytical estimate of the response time. We created the data in a) by scanning *n*_RF_, setting 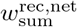 to yield a RF of size *n*_RF_, setting 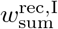 to 0 or its critical value, and determining *τ*_resp_ or 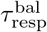 from the loss dynamics. For the SFA network we set *τ*_SFA_ = *τ*, scanned *a*_SFA_ and used the value that minimized the temporally integrated loss.

### Linear mixed selectivity

Neurons often respond selectively to more than one stimulus or input feature [64]. This phenomenon is called mixed selectivity (MS). Here we consider linear MS [64, 65], where neuronal responses are linear functions of multiple stimuli. Concretely, neurons with activities *x*_*ij*_, *i, j* = 1, …, *N* are arranged on a two-dimensional grid and respond with equal selectivity to two input features, represented by input neurons 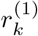 and 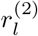 with *k, l* = 1, · · ·, *N*,

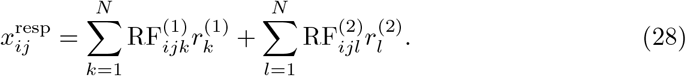

Due to the linearity in the input representation and in the network, the total response is the sum of the responses to the single input features. We take the grid axes to be aligned with the stimulus dimensions, so that the first index in *x*_*ij*_ determines its response to *r*^(1)^ and the second that to *r*^(2)^. We model this dependence as the same localized, exponentially decaying shape as for the 1D network (cf. Fig. 6b)),

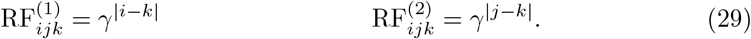

the desired network response can be generated as the steady state of a recurrent network that is equivalent to the 1D network eq. (5) in each dimension of the 2D grid (see next section),

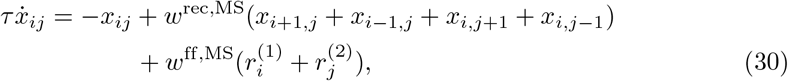

with the modified constants 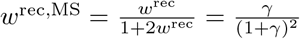 and 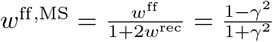. Each neuron receives two external inputs and is connected to its nearest neighbors along each stimulus axis. The network has thus only six synapses per neuron, regardless of the RF width. Also a feedforward network where each dimension of the 2D grid is equal to the 1D network eq. (2) generates the desired response. This implementation requires 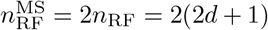 synapses per neuron, a number that increases linearly with the RF width.

### Mapping to a 1D system

In the following, we trace the network dynamics eq. (30) back to those of the 1D system eq. (5). Due to the linearity of eq. (30), network responses again superpose. It thus suffices to study the network in the case where only one input neuron is active: we choose 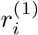, which specifies a property of the first stimulus, to be nonzero. Since the input is independent of *j*, the dynamics eq. (30) are (for initial conditions homogeneous in *j* such as *x*_*ij*_(0) = 0) independent of *j, x*_*ij*_(*t*) = *x*_*i*_(*t*). The recurrent inputs *w*^rec,MS^(*x*_*i*,*j*+1_(*t*) + *x*_*i*,*j*−1_(*t*)) = 2*w*^rec,MS^*x*_*i*_(*t*) then simply amount to a modification of the leak current to −(1 − 2*w*^rec,MS^)*x*_*i*_,

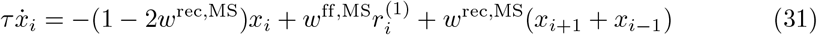

After dividing by (1 − 2*w*^rec,MS^), the differential equation for *x*_*i*_ becomes

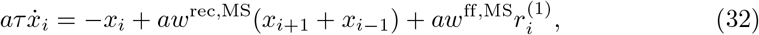

where we introduced *a* = (1 − 2*w*^rec,MS^)^−1^ for brevity. With the values of the constants *w*^rec,MS^ and *w*^ff,MS^ highlighted after eq. (30) this is equivalent to the one-dimensional network dynamics eq. (5) up to a different neuronal time constant *aτ* instead of *τ*. (We note that we obtained the modified constants such that this holds. For example, equating the prefactors of the recurrent term in eq. (32) and eq. (5) gives *w*^rec^ = *aw*^rec,MS^ = (1 − 2*w*^rec,MS^)^−1^*w*^rec,MS^, which then can be solved for *w*^rec,MS^.) As a direct consequence, while the 1D network must have recurrent coupling strength of *w*^rec^ *<* 0.5 for being stable, the 2D MS network must have *w*^rec,MS^ *<* 0.25. This is because in the MS network, a neuron receives direct recurrent input from four nearest neighbors instead of two as in the 1D case.

Equation (32) means that the RF of the MS network, along one axis, has the same shape and width *d* as the equivalent one-dimensional network. In particular, *d* is related to the recurrent weight strength 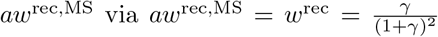 and *γ* = exp(−1*/d*); the two RF components in eq. (29) are the same as the RFs in eq. (1), for example 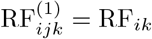.

### Response speed

From the mapping of the MS to the 1D system, eq. (32), we see that the MS dynamics behaves in response to a single input like the 1D dynamics with the neuronal time constant *τ* enlarged by a factor of *a*. The response time is thus given by eq. (12), but with enlarged neuronal time constant, *τ* → *aτ*. For sufficiently large *d*, we have *γ* ≈ 1 (reflecting the spatially slow RF decay), *w*^rec^ ≈ 1*/*2, *w*^rec,MS^ ≈ 1*/*4 and thus *a* ≈ 2. The scaling of the response time with the RF width *d* and size 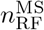 is thus again quadratic,

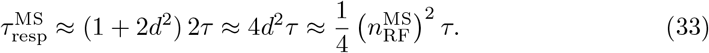

In the last equation we used 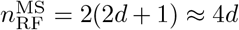. Compared to the 1D case (eq. (12)), the response time as a function of *d* is therefore larger by a factor *a* ≈ 2. In contrast, it is smaller by a factor 1*/*2 as a function of the RF size, compare eq. (33) with eq. (12) and the blue continuous and dotted curves in Fig. 6a). In other words: the trade-off between response time and number of needed synapses improves for sufficiently large RFs by a constant factor of about 1*/*2 compared to the 1D network. This is because the MS network effectively implements two 1D RFs (eq. (28)).

### Balanced network

We now incorporate the effect of inhibitory neurons into the MS network. As in the 1D case, we assume that the generated inhibition precisely tracks excitation with a short time delay. We thus add to each recurrent excitatory connection an inhibitory one that is slightly delayed. This results in a delayed differential equation like eq. (24) for the balanced MS network dynamics. The parameters are given by those of the 1D balanced system up to a factor 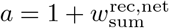, like in eq. (32). Further, it is again sufficient to study the response dynamics to a single input, which can be reduced to those of the 1D balanced network eq. (24) with adapted parameters. As in the purely excitatory case, for a fair comparison of response times, we consider MS and 1D networks with the same neuronal time constant *τ*. The effective time constant of the MS dynamics are then *aτ*. Therefore the response time of the MS network are given by those of the 1D network eq. (27) with neuronal time constant *τ* replaced by *aτ* ≈ 2*τ*,

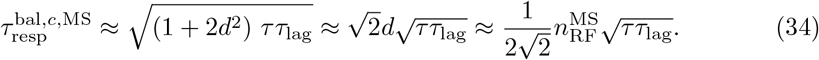

At the critical inhibitory strength, the response time thus scales again with the square root of the response time of the network without inhibition (eq. (26)). Therefore it scales linearly with the receptive field width *d* and size 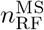. The response time is as a function of the RF size by a factor of about 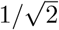 smaller than that in the 1D case, eq. (27), see Fig. 6a), red continuous and dotted curves. In other words, the trade-off between response time and number of required synapses improves by a factor of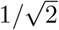. This is again because the MS network effectively implements two RFs in the MS case; the RF size doubles for the same width compared to the 1D network.

As for the 1D stimulus, the balanced networks have twice as many neurons and additional synapses: Each (excitatory) feature neuron drives one inhibitory neuron, which mirrors its activity. This inhibitory neuron in turn forms inhibitory synapses to the four nearest neighbors that its presynaptic feature neuron excites. In total, the balanced, cooperatively coding network thus requires eleven synapses per feature neuron, instead of six for the unbalanced network. This is independent of the RF width, such that the balanced, cooperatively coding network save synapses for sufficiently wide RFs.

### Higher-dimensional linear MS

We can straightforwardly extend the introduced scheme to networks that have MS with *P >* 2 stimuli. Neurons are then arranged on a hyper-grid with one grid axis per stimulus dimension, so that *N*^*P*^ feature neurons respond to *PN* input neurons. In the cooperatively coding network, each neuron receives *P* feedforward and 2*P* recurrent inputs, requiring a total of 3*P* synapses per neuron. The feedforward network, in contrast, needs for each stimulus dimension 2*d* + 1 synapses, in total *P* (2*d* + 1) synapses per neuron. The number of saved synapses thus grows linearly with the number of encoded stimulus dimensions and the receptive field width.

## Encoding a 2D stimulus

We finally consider the encoding of a two-dimensional stimulus, with both input and feature neurons arranged on a two-dimensional grid, see Fig. 7a). Two-dimensional input appears for example in vision [36] or planar navigation tasks [66]. Each feature neuron responds to inputs that are close to its preferred input in both stimulus dimensions. The RFs of neighboring neurons thus overlap and neuronal responses tile the represented stimulus space. A purely excitatory cooperative coding network generating such activity as stationary state is given by

**Fig 7:**
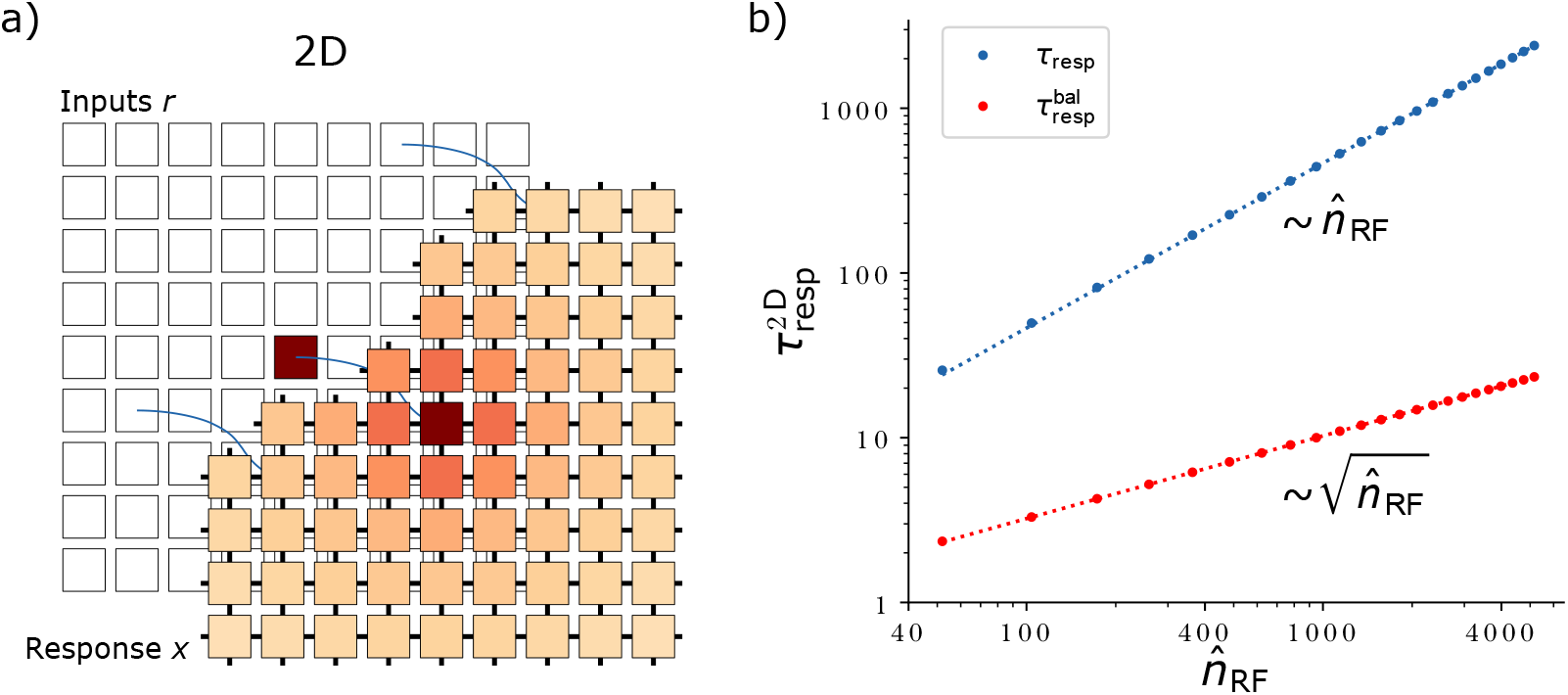
2D network schematic and response times versus RF size. **a)**Schematic of a two-dimensional network responding to a two-dimensional stimulus. Feature neurons (labeled ‘Response *x*’) and input neurons (labeled ‘Inputs *r*’) are arranged on two-dimensional grids. In the cooperatively coding network each feature neuron receives one feedforward and four recurrent inputs; activities and shown connections are color-coded as in Fig. 2. **b)** Response times in the 2D network increase linearly (without inhibition, blue. Monomial fit: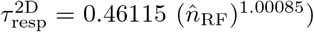) or square-root-like (with inhibition, red. Monomial fit: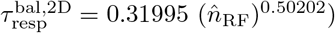) with the RF size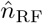. Dotted lines represent the monomial fits. Data was created by scanning 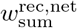, setting 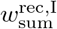 to 0 or its critical value, and determining 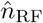 and *τ*_resp_ or 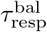, respectively, from the response curves after network activity converged.

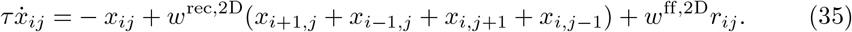

It has the same recurrent connectivity as the network with linear MS eq. (30), but the feedforward input is arranged on a grid. The activity of feature neuron ij in the stationary state is

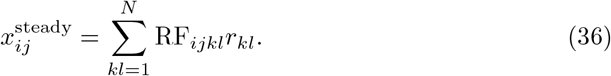

the neuron thus responds to a combination of two input features represented by input neurons *r*_*ij*_ with *i, j* = 1, *· · ·, N*. The RF is explicitly given by

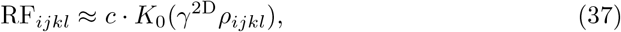

with 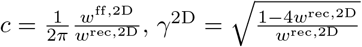 and 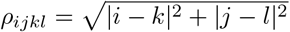, *K*_0_ is the zeroth modified Bessel function of the second kind, which decays with distance ρ approximately as 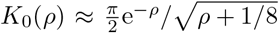 [67]. The RF is thus approximately radially symmetric. The RF size depends on *w*^rec,2D^ and the response amplitude also on *w*^ff,2D^. In contrast to the 1D and the MS case, these dependencies cannot easily be computed analytically; in Fig. 7 we thus determine the receptive field sizes resulting from the tried *w*^rec,2D^ numerically. The network requires 5 synapses per neuron.

We also constructed a balanced network by introducing for each excitatory recurrent input a delayed inhibitory one, like in the balanced 1D and MS networks. A balanced implementation with explicit inhibitory interneurons requires twice as many neurons and synapses than the purely excitatory network: it requires additionally one interneuron per principle neuron, one inhibitory synapse for each excitatory recurrent synapse and one synapse from each principle neuron to its corresponding interneuron.

### Response speed

To estimate the dependence of the response speed on the RF size, we first need to appropriately adapt the definition of the RF size, which we introduced after eq. (3). For the one-dimensional network, this definition can be reformulated as follows: we count the number of synapses that are necessary to generate the largest (around the center) RF responses such that these responses summed together amount to a fraction of about 1 − *e*^−1^ ≈ 63% of the summed non-truncated RF. Accordingly, for the 2D network at hand we define the RF size as the number of feedforward synapses that are necessary to implement the largest RF entries, such that together they account for a fraction of approximately 63% of the summed nontruncated RF. We denote the so-defined receptive field sizes by 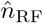.

As for the 1D and linear MS networks, the response time with or without lagged inhibition depends only on the summed excitatory weights or on the summed net and inhibitory weights. It is thus given by eq. (11) or by eq. (26) in terms of 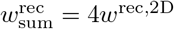 or in terms of the (alike obtained) 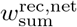 and 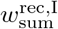. Fig. 7b) shows that the scaling of the response time with the RF size is linear for unbalanced and square-root-like for balanced networks. We give a geometric argument for this general scaling in the Discussion (subsection ‘Multi-dimensional stimuli’). The scaling is more economical than for the 1D and 2D linear MS networks, cf. Fig. 6.

## Discussion

In this work, we have studied networks that encode continuous variables with neurons that have overlapping response properties. We developed a cooperative coding scheme that enables them to share and distribute computations among similarly-tuned neurons, crucially using (net) excitatory connections. For the simplest considered networks this sharing minimizes the number of required synapses while the total amount of synaptic current remains the same as in a purely feedforward implementation. For networks of neurons that represent higher-dimensional stimuli, the number of saved synapses is especially large. The saving of synapses comes at the cost of longer response times. We find, however, that neurons with spike frequency adaptation and neurons in networks in which excitation is largely balanced by delayed inhibition can use the window of opportunity between the arrival of excitation and inhibition to significantly speed up their convergence to the steady state response, decreasing response times by orders of magnitude and improving their scaling with RF size.

### RF shape and more complex networks

The exponentially decaying RFs that we consider act as tractable models for experimentally encountered localized, overlapping and broadening RFs [2, 38, 40]. They allow to elegantly illustrate how neurons can use recurrent interactions to cooperatively share feedforward information and shape the network response. However, experimentally measured RFs have different and often more complex shapes [7–9, 36]. We are optimistic that these can still be approximated by the steady state of a cooperatively coding recurrent network with sparse connectivity, although more synapses will be required. Determining the necessary network parameters might involve minimizing a loss with L0 regularization, which is challenging.

### Related work

Conceptually, our 1D model is a ring model. Models of this type have been proposed to model orientation selectivity in the visual cortex [23, 52] (see also [68]), head direction cells [69] and spatial memory [70]. Similarly, our 2D and 2D mixed selectivity models have a toroidal or, when removing the periodicity, a planar structure. Such networks may be important for spatial navigation [66]. With appropriate coupling, ring-like networks can have two different dynamical regimes [42]: (i) an input-driven regime where there is a single, homogeneous ground state, which is assumed in absence of input and (ii) a regime of bump attractors, where there is spatially localized, persistent activity in absence of input. Our networks are linear, therefore without input there cannot be multiple stationary activity patterns whose amplitudes are asymptotically stable. This is because each multiple of a stationary solution is a solution as well. We thus work in the input-driven regime, with a single stable zero ground state. Previous models have broad coupling fields or ranges of coupling probabilities, equivalent to many recurrent synaptic connections that extend over neurons with quite different preferred stimuli [23, 42, 52, 66, 69, 70]. In contrast, in cooperative coding networks, we have very sparse synaptic connections between neurons with highly similar tuning.

In this work we considered the encoding of continuous variables in a scheme that minimizes the total number of required synapses. Relatedly, ref. [71] investigates the problem of classifying discrete patterns in large networks of neurons with limited and fixed in-degrees. It solves the problem by introducing an intermediate layer consisting of interconnected perceptrons, which each receive only part of the overall input. This intermediate layer may be seen as analogous to our feature layer. In the mainly studied binary classification task, the intermediate layer is equipped with excitatory connections. The assumed connectivity is untuned except for being overcritically excitatory such that it drives neurons into saturation. In the stationary response to a pattern then all (more precisely: nearly all [72]) neurons in the network assume the same output, either positive or negative one. This allows a readout neuron to perform binary classification with sparse readout weights. A refined architecture combines several such intermediate layers in parallel. This yields an intermediate layer with several subpopulations, which code in binary manner, allowing for classification with a binary vector. Another way highlighted to achieve this is by using as intermediate layer a Hopfield network with recurrent connections that are randomly realized with fixed probability. Since the binary coding network and the network with several subpopulations have specific like-to-like connectivity to save feedforward connections, they realize cooperative coding in our sense, in a binary manner. By choosing the recurrent coupling probability of the Hopfield network intermediate layer such that stronger positive and negative weights (which connect similarly and oppositely tuned neurons, respectively) are realized with larger probability, one could implement cooperative coding in them as well. Ref. [73] find that local recurrent connectivity in Hebbian assemblies of spiking neurons can reduce the number of feedforward connections between assemblies required for memory replay. The total number of synapses in their model is, however, minimized by a purely feedforward architecture. They argue that one benefit of sparse feedforward connectivity, augmented through local recurrence, might be enabling one-shot learning.

The signature of cooperative coding in the networks that we investigated and in more general ones is that the network trades feedforward and less specific recurrent synapses for recurrent synapses mediating on average net-excitatory recurrent interactions between similarly tuned neurons (and net-inhibitory interactions between highly anti-tuned neurons). Consistent with this, intermediate-depth ML networks featuring recurrent and feedback connections can match the performance of much deeper feedforward networks while requiring less units and parameters [74]. It would be interesting to investigate whether the recurrent connectivity in such networks is also like-to-like. If yes, this would indicate that cooperative coding naturally appears also in ML networks. It may be helpful in particular in convolutional networks to save feedforward connections and rely on very sparse recurrent connectivity instead.

### Response speed in excitatory 1D networks

In our most simple, purely excitatory cooperatively coding networks the response is slowed down compared to that of single neurons due to recurrent excitation, which implements a positive feedback loop. This feedback loop increases the eigenvalues of the matrix governing the differential equation of the dynamics and increases the duration of the response to single pulses. The prolonged response to a single short pulse adds up for prolonged inputs and leads to their amplification as well. This type of amplification has been termed “Hebbian amplification” in [21].

### Networks with SFA

To speed up the response, we first introduced SFA, a typical feature of excitatory principal neurons [42, 49, 50]. Our model is a slightly simplified version of that in [51] and the same as in [52, 53].

In the networks with SFA excitation still dominates, i.e. we have on the one hand Hebbian amplification as well. On the other hand, we have “balanced amplification” [21]: Each neuron is a 2-dimensional dynamical system, the network consists of *N* of them; they are excitatorily coupled. Without the inhibitory feedback from the adaptation the dynamics of the purely excitatory neuron population would be unstable. When a short input arrives to the neuron population, its activity therefore increases rapidly, using the window of opportunity before the inhibitory feedback current becomes prominent. Inhibition then takes over and largely terminates the excursion leading to a shorter duration impulse response of the network. For prolonged input, these impulses basically add up, which leads to the fast convergence to new stationary activity, amplifying the input. The complex Schur decomposition of the matrix governing the network dynamics reveals a strong feedforward coupling from an oscillatory difference to an oscillatory sum mode. This is similar to the networks in [21], which, however, have mostly real, non-positive eigenvalues and thus non-oscillatory modes without Hebbian amplification.

### Balanced networks

The balanced versions of our networks incorporate inhibition that closely tracks excitation with a short delay, as observed in experiments [60, 75, 76]. For stationary dynamics, excitatory and inhibitory currents largely cancel, resulting in a relatively weak net excitatory interaction between similarly-tuned neurons [18, 19]. However, excitation reacts slightly faster than inhibition to dynamic activity changes, allowing strong interactions during the window of opportunity where the change in excitation is not yet balanced by the delayed inhibition. This mechanism allows activity changes to propagate quickly through the network, significantly decreasing its response time.

The effective lag of inhibitory feedback originates in our SFA and in other balanced amplification [21] networks from the fact that the inhibitory currents are evoked by a low-pass filtered version of the excitatory activity. In contrast, in our balanced networks, there is an explicit, fixed lag between excitatory and inhibitory activity. We thus have an infinite dimensional dynamical system governed by a delay differential equation. However, the basic mechanism of shortening the impulse response and speeding up the reaction to inputs is the same: Excitation without feedback inhibition is unstable. When an input arrives, inhibition is insufficient to balance it. The excitatory activity therefore increases strongly during the temporal window of opportunity. Then inhibition sets in, largely balances excitation and limits it to its stationary value. We note that still excitation dominates our balanced networks. We have numerically and analytically studied the resulting convergence to stationary activity. This revealed qualitatively different types of dynamics, which are familiar from the harmonic oscillator, namely overdamped, critical and underdamped dynamics. Their occurrence depends on the strength of the inhibitory feedback and the size of the time lag between excitation and inhibition.

The increased response speed comes at the cost of requiring additional synapses and interneurons to implement the inhibition. In our model this roughly doubles the number of required synapses compared to the unbalanced cooperatively coding networks. As the number of needed synapses is still independent of the RF size, the balanced networks still save synapses for larger RFs. Furthermore, inhibitory neurons may be required anyways for purposes such as maintaining irregular spiking activity [55–58]. Alongside the increased synaptic requirements, the cancellation of currents means an increased metabolic cost.

We covered inhibition in a simplified, effective manner without explicitly incorporating inhibitory neurons. As argued above, we are optimistic that our qualitative findings will hold in more complex networks, for example if they are optimized for sparseness. In particular we expect that such networks will still show characteristics of cooperative coding and that inhibition will allow to reduce response times.

From our analysis, the brain might use cooperative coding to save synapses and space compared to a purely feedforward or more wasteful recurrent implementation, but might invest some synapses, neurons, space and energy in balancing inhibition to retain a reasonable response speed.

### Mixed selectivity

We also considered networks with two-dimensional mixed selectivity, where neurons are not only selective to a single stimulus but to two stimuli. Since our networks are linear, we consider linear mixed selectivity. This is a simplification compared to the nonlinear mixed selectivity that is ubiquitous in the brain [64, 65]. We observe that the tradeoff between saved synapses and speed is for networks using linear mixed selectivity improved by a constant factor compared to networks where each neuron codes for a single stimulus.

### Multi-dimensional stimuli

We considered networks encoding one- and two-dimensional stimuli. We found that the trade-off between saved synapses and increased response time is much more favorable for neurons that encode 2D stimuli.

For *P* ≥ 2-dimensional stimuli, the fact that activity simultaneously propagates along all dimensions suggests that the scaling of the response time with *d*, the characteristic RF width along one of the dimensions, does not change with the number of dimensions. However, the RF size *n*_RF_, the number of synapses needed in a purely feedforward implementation, can be assumed to scale as *n*_RF_ ∼ (2*d* + 1)^*P*^, with a prefactor that depends on geometry. This reasoning suggests that the response time of networks encoding higher-dimensional stimuli scales like 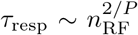 and 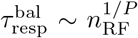, which we verified for *P* = 1, 2. The number of synapses required in a cooperatively coding network will also in higher dimensions be negligible compared to *n*_RF_. Therefore the number of saved synapses in the cooperatively coding network is still approximately *n*_RF_. For higher-dimensional stimuli, the trade-off between response time and saved synapses would thus become highly beneficial: the response time *τ*_resp_ or 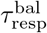 would grow only slowly with the number of saved synapses *n*_RF_ due to the strongly sublinear relationship for larger *P*.

### Properties of connectivity

Our cooperative coding scheme relies on the presence of few strong recurrent excitatory connections between similarly-tuned cells. Excitation can be balanced by inhibition, while interactions between similarly-tuned cells remain net excitatory. Indeed, in layer 2/3 of mouse visual cortex, pyramidal neurons with the same orientation preference connect at higher rates and form more bidirectional connections [15]. This pattern of increased connection probability between neurons with highly similar tuning extends across layers and visual areas, including feedforward and feedback connections [14]. Furthermore, synapses between neurons with similar spatial RFs are markedly stronger such that neurons receive the majority of their local excitation from few similarly-tuned cells [16]. This strong, sparse local excitation matches the RF structure of the receiving neurons [16]. Recurrent connections are generally sparse in the cortex [77–80].

Studies of recurrent functional connectivity found net excitation between (spatially close) neurons with similar tuning [19] and most correlated responses [18], consistent with our model. Ref. [19] showed specifically that when optogenetically stimulating spatially compact ensembles of co-tuned neurons, similarly-tuned neurons were excited while differently-tuned neurons were inhibited. In a 1D-model, ref. [18] had to incorporate strong nearest-neighbor-like excitatory interactions to match the experimental data, arguing that they stabilize network responses in the presence of input noise. Our interpretation is that they not only stabilize, but actively form the RF. Refs. [18, 19] also show an inhibitory effect on largely differently tuned neurons. Ref. [18] found net inhibition between rather similarly tuned neurons as well. This is assumed to implement feature competition, which we did not include in our model.

A particular benefit of our cooperative coding scheme is that it allows feedforward connections to be sparse. This fits for example experimental observations in the primary visual cortex, where the vast majority of inputs are local recurrent ones, while only a few percent are feedforward inputs [81, 82]. Ref. [83] estimated based on experimental studies [84] that a single hypercolumn in primate V1 receives only 10-30 feedforward inputs from the magnocellular layer of dorsal LGN mediating retinal input, with single cells in L4*α* receiving as little as 0 - 6 inputs. Also in the hippocampal region CA3, where place fields are enlarged compared to the upstream dentate gyrus [40], the recurrent connectivity is high [79, 85].

Experiments that aim to disentangle feedforward from recurrent contributions to orientation selectivity resulted in mixed findings. Ref. [86] showed that the input to simple cells in L4 of cat primary visual cortex still exhibit tuned, modulated responses to drifting gratings after cortical activity was suppressed by cooling. In line with this, ref. [17] found that thalamic and cortical contributions to the first harmonic of the response curve (F1) were co-tuned. However, the temporally averaged response (F0) to drifting gratings was tuned only in cortical but not in thalamic inputs. A recent study, ref. [87], suggests that total input current from L4 of mouse primary visual cortex to L2/3 may lack orientation tuning and that orientation selectivity is determined by recurrent inputs from within L2/3.

### Optimality

By distributing and reusing computations via excitatory connections, cooperatively coding networks minimize or strongly reduce the number of synapses required to generate their responses. This corresponds to minimizing or strongly reducing the L0 norm of the weights (or synaptic currents).

In the following we compare the metabolic cost of the feedforward and the simplest, purely excitatory cooperative coding networks in the stationary state. We consider the three main contributions of the metabolic cost [26, 28]: the cost of keeping up the resting network, of generating the required activity and of the synaptic transmissions. Keeping up the resting network (housekeeping and maintaining the resting potentials) requires the same energy expenditure in both implementations, as the number of neurons is the same. Also the energy required to generate the stationary output activity is the same, as corresponding neurons generate the same activity in both implementations.

The cost of synaptic transmission is dominated by the metabolic cost caused by the postsynaptic currents [28]. We assume that this cost is characterized by the sum of the absolute current strengths at individual synapses, i.e. by the L1 norm of the synaptic currents [88]. This is the same in both implementations. Finally, a small part of the overall cost (less than 10% [28]) arises due to presynaptic calcium influx and usage of neurotransmitter during synaptic transmission. A comparison of these contributions between the network implementations is more difficult. The surface of the active zone [89] increases linearly with the synaptic strength [90]. Assuming that the area where calcium influx happens increases linearly with the active zone size, the amount of influx during a single synaptic transmission depends linearly on the synaptic strength, consistent with [91]. The same should then hold for the related cost. Concerning the amount of neurotransmitter used, experiments observe that the number of directly releasable vesicles is linearly related to the synaptic strength, while the impact of a single vesicle and its release probability are independent of it [90, 92] (this may differ for different connected neuron types [93]). The finding suggests a linear dependence between the synaptic strength and the used neurotransmitter and thus the related cost at a single transmission. The total cost due to presynaptic calcium influx and neurotransmitter usage is therefore proportional to the absolute synaptic strength times the number of transmissions (the presynaptic activity). This, in turn, is proportional to the absolute value of the induced postsynaptic current, which is identical in both implementations.

We conclude that the metabolic cost of generating the stationary state is the same in the feedforward and the simplest, purely excitatory cooperative coding implementation. Since the latter takes longer to reach the stationary state, however, it also requires more current and energy before generating a useful result. Due to the cancellation of excitatory and inhibitory currents, the networks with SFA and the balanced networks also use more current and thus more energy in the stationary state.

Previous studies often minimized the number of spikes or, more generally, the neuronal activity needed to represent encoded features. Refs. [30, 32, 34] follow this approach and suggest that tight EI-balance may be a signature of a highly coordinated and competitive code that, despite the irregular firing, is orders of magnitude more precise than a Poisson rate code. This spike-code depends on an extremely structured, dense connectivity, through which similarly-coding neurons quickly inhibit each other to prevent redundant spiking. From this standpoint the findings of excitatory functional connectivity between similarly-tuned neurons [15, 16, 18, 19] seem counter-intuitive.

Although the energetic costs of spike generation and synaptic transmission very likely play a role in shaping cortical networks, our work suggests that space constraints and the number of required synapses may be another main factor.

### Conclusion

To conclude, net excitatory connectivity between similarly tuned neurons is compatible with a cooperative coding scheme that generates network responses with a minimal number of synapses. This suggests space constraints as an important factor in shaping neural networks. The window of opportunity between excitation and balancing, delayed adaptation or inhibition may be harnessed to rapidly propagate activity changes through the network, speeding up equilibration times by orders of magnitude.

## Methods

All simulations have periodic boundary conditions. Fixed network parameters are the number of neurons *N* for 1D and *N* ^2^ for 2D networks, the neuronal time-constant *τ* and, in networks with inhibition, the EI-lag *τ*_lag_. We set *N* = 200, *τ* = 1 and *τ*_lag_ = 0.1. In the networks with SFA we use a fixed value of *τ*_SFA_ = *τ* = 1 and, for each RF size, obtain the value of *a*_SFA_ that minimizes the temporal mean of the normalized L1-loss, 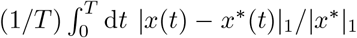, through a linear grid search. Here *T* = 500*τ* is the length of a trial as described in Fig. 8a) and *x*^∗^(*t*) the target corresponding to the present input. Fig. 8e) and c) show the scans over *a*_SFA_ and the individual loss curves for the optimal *a*_SFA_ values. In all networks with inhibition, we set 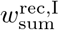 to its critical value given by eq. (77).

**Fig 8:**
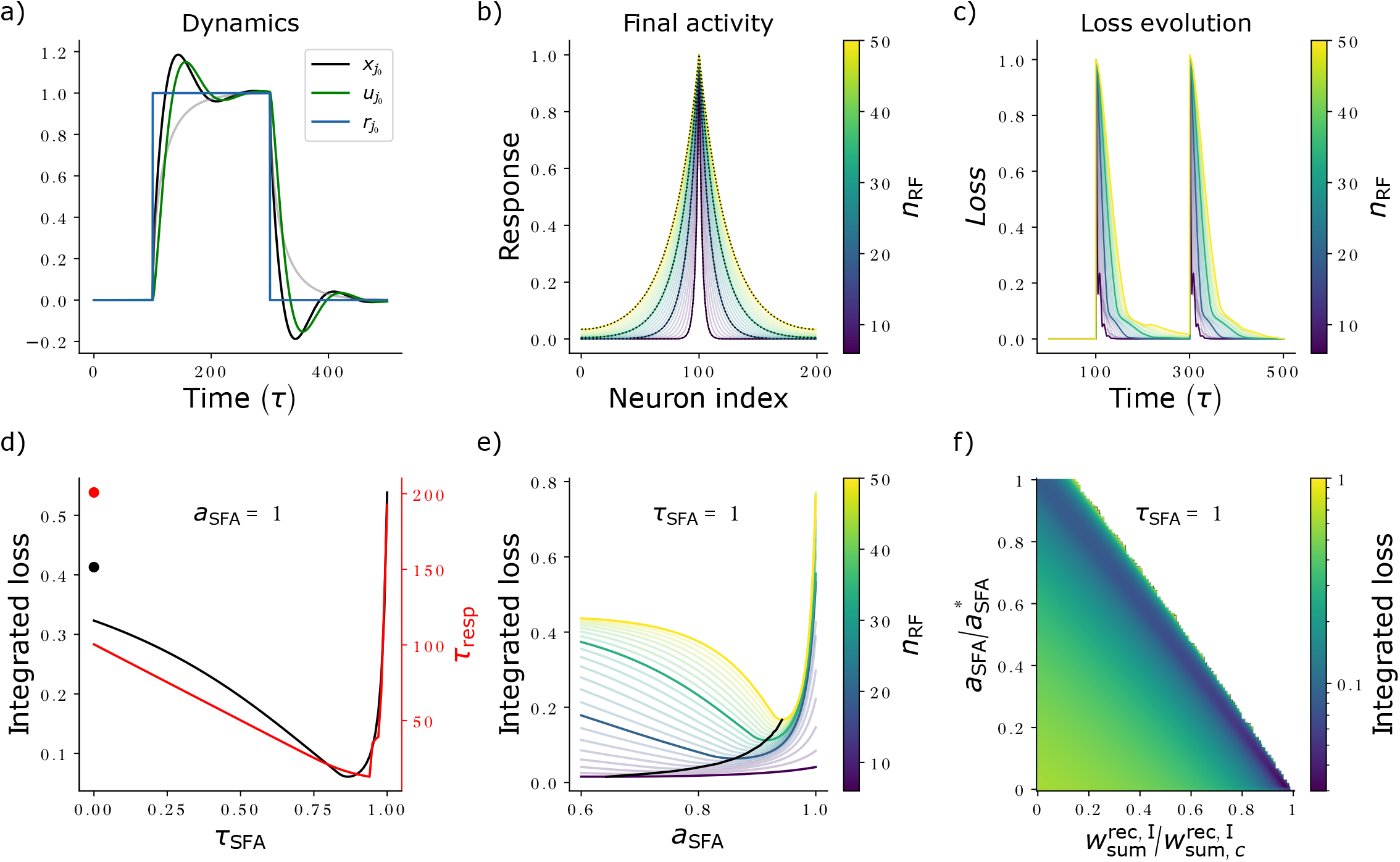
Network and loss dynamics in a 1D network with SFA. **a)** Dynamics of activity (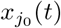, black), adaptation variable (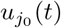, green) and input (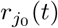, blue) for the *j*_0_ = 100th neuron, which receives its preferred input as isolated unit input that is switched on at *t* = 100*τ* and off at 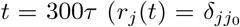 between these times). The network implements an RF with *d* = 4.5 (*n*_RF_ = 10) and has, for better illustration, a rather slow *τ*_SFA_ = 10*τ* and slightly stronger-than-optimal *a*_SFA_ = 0.09, resulting in a visible lag between *x*(*t*) and *u*(*t*) and oscillatory dynamics. During the initial rising phase (*t* ≳ 100*τ*), activity rises faster than for a network without SFA (gray line). **b)** Stationary activity of SFA networks with different receptive field width. The networks receive an isolated input (parameters are *τ*_SFA_ = *τ* and optimal 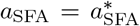, see e)). The stationary activity matches its target for different *n*_RF_ (color-coded). To show this, target (black dashed) and final activity (color-coded) of four networks with *n*_RF_ 6, 20, 34, 50 are highlighted. **c)** Evolution of the L1-loss of the deviation of network activity from its target, normalized by the L1-norm of the target response for present input, under the stimulus protocol as described in a) for the same networks as in b), with *a*_SFA_ determined as described in e). **d)** Integrated loss (black), determined as the temporal mean of the normalized L1-loss shown in c), and response time (red), determined as the earliest time after the onset of input for which the normalized L1-loss drops and stays below e^−1^, for a network with *d*_RF_ = 10 (*n*_RF_ = 21), *a*_SFA_ = 1 and 101 values of *τ*_SFA_ scanned between 0 and 1. The integrated loss and response time of a network without SFA are shown by colored dots. **e)** Integrated loss for different *n*_RF_ (curves are color coded as in b)) as a function of *a*_SFA_ (for *τ*_SFA_ = 1). The minima, determining 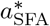s used in b), c) and Fig. 6a), are connected by a black curve, showing that the optimal adaptation strength increases for larger RFs. We note that the data in a),d) and e) suggest that the optimal adaptation parameters fulfill the relation *a*_SFA_*τ*_SFA_ ≲ 1.f) Grid scan of the integrated loss for *a*_SFA_ linearly scanned between 0 and 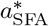 and 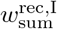 linearly scanned between 0 and 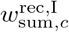 (101 values each, *d*_RF_ = 10 (*n*_RF_ = 21), *τ*_SFA_ = 1). The white region indicates parameters for which network activity diverges, identified by an integrated loss larger than 1, cf. also e). Here the loss was computed for the time window with present input only, more precisely for *t* ∈ [100*τ* − d*t*, 300*τ*], different from a)-e). For all simulations, we created data by simulating 1D networks with *N* = 200, *τ* = 1 using the Euler method with step size d*t* = 0.01; *n*_RF_, *τ*_SFA_, *a*_SFA_ and 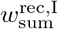 varied as described above.

**Fig 9:**
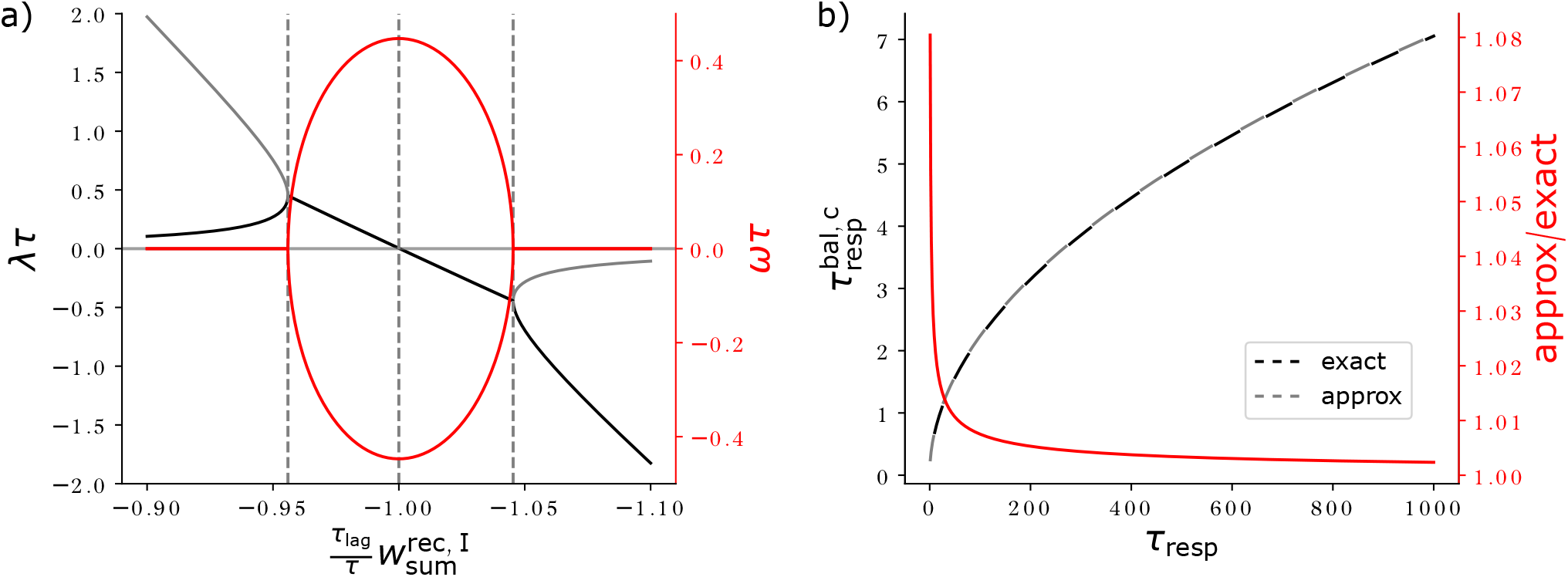
Critical points and approximation of the analytical solution for 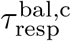. **a)** Real part (decay rate *λ*, black/gray) and imaginary part (oscillation frequency *ω* times ± 1, red) of the complex frequency of the exponential loss evolution, scaled by *τ* (compare Fig. 5). The leftmost dashed gray vertical line marks the first critical inhibitory strength at which there is only a single, positive decay rate. For 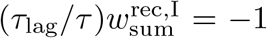, marked by the middle dashed gray vertical line, the decay rate is zero and transitions from positive (decaying) to negative (exponentially growing). There is a second critical inhibitory strength, marked by the rightmost dashed gray vertical line, at which there is a single, negative decay rate. The critical point with the decaying dynamics corresponds to the solution of eq. (77) with *k* = 0, the one with the exponentially growing dynamics to that with *k* = − 1. **b)** Exact (black dashed, cf. eq. (80)) and approximate (gray dashed, cf. eq. (89)) values of the critical decay time of the balanced network, 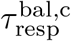, as a function of that of the excitatory network. Approximation and exact solution agree well. Their ratio (red) is close to one, and approaches one for *τ*_resp_ ≫ *τ*_lag_. Parameters: *τ* = 1, *τ*_lag_ = 0.1 and, in a), 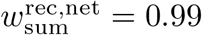.

We simulate our networks with SFA using the Euler method and all other networks using the midpoint method with stepsize d*t* = 0.01. To simulate the networks with delayed inhibition, we also need midpoint values of the delayed activity. We obtain them by copying the midpoint values of the non-delayed activity *τ*_lag_ (*τ*_lag_*/* d*t* simulation steps) before.

For the data in Figs. 4 and 6, we obtain RFs with different sizes by setting 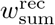 or 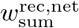 to appropriate values. In the case of 1D networks with and without SFA and in the case of 2D linear MS networks, we have analytical expressions for the RF sizes as a function of 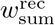 or 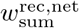. We thus chose 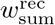 or 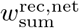 such that the RF sizes are sampled linearly from 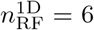 to 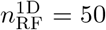 in steps of two. For the 2D network, we simulate networks with 20 different values of 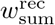 or 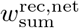 and measure the RF sizes that the networks generate after convergence. We obtain 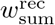 or 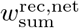 as 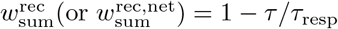 by varying *τ*_resp_ from 10 to 1, 000 with equal spacing on a logarithmic scale.

To numerically determine a network’s response time, we first simulate the network for a long time, clearly longer than the convergence time, and define the resulting state as the final, target state *x*^∗^. The loss is the L1 norm of the difference between *x*^∗^ and the current state. We then simulate the network for a second time. We obtain *τ*_*resp*_ or 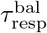 as the earliest time at which the loss drops and stays below e^−1^ times the initial loss.

## Acknowledgments

We thank Felipe Kalle Kossio for fruitful discussions.

## Supporting information

### S1 Dynamics of excitatory networks

The dynamics of our networks without SFA and explicit inhibition read for constant input *r*_*i*_(*t*) = *r*_*i*_

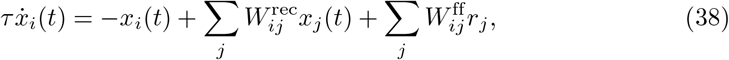

see eq. (4). In the following, we discuss stationary states and time-dependent behaviors of these dynamics.

We first verify that the recurrent network eq. (5), where 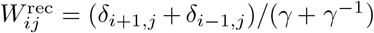 and 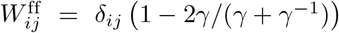, has the desired stationary state 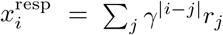. For this we insert eq. (1) into eq. (5),

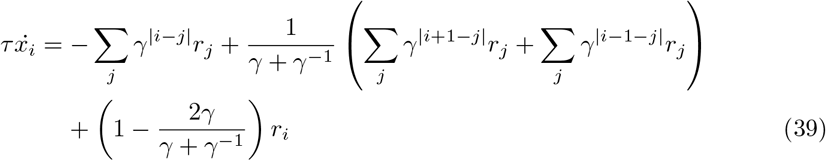

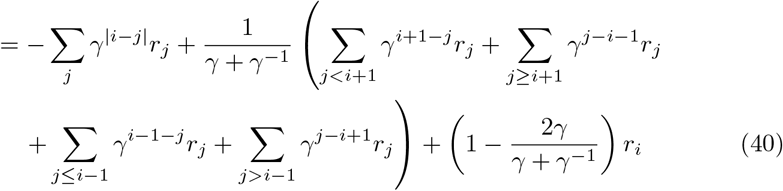

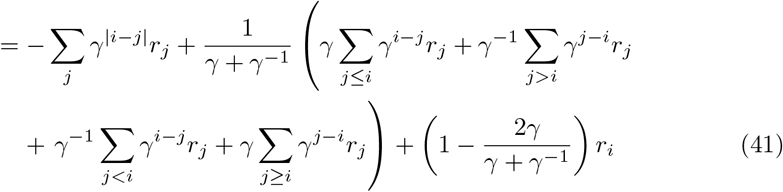

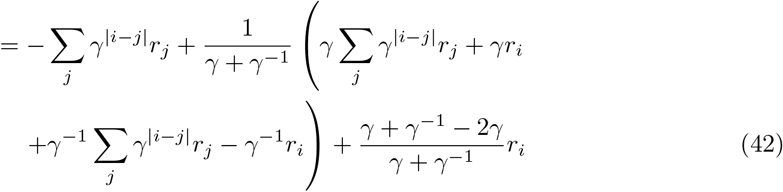

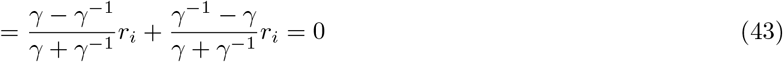

the computation shows that the recurrent input from neurons *i* + 1 and *i* − 1 add, in the steady state, up to nearly generate the desired response of neuron *i*: the first and third summand in the bracket in eq. (42) give the desired response. The input that is missing,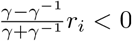, is contributed by the feedforward input (last summand in each line, which cancels the missing input in eq. (43)).

We now turn to non-stationary solutions. The recurrent weight matrices in our recurrent networks eq. (5) are real symmetric matrices and therefore diagonalizable with real eigenvalues and orthogonal eigenvectors. We denote by 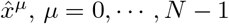, the *μ*th normalized eigenvector of *W* ^rec^ with eigenvalue *α*^*μ*^, and by 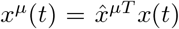 the projection of the network activity on this eigenvector (a scalar). Any activity *x*(*t*) can be expressed as a linear combination of the orthonormal eigenvectors 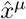 with coefficients 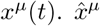 is also an eigenvector of *W* ^rec^ − 𝟙, with eigenvalue *α*^*μ*^ − 1. Multiplying eq. (38) with 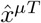 from the left shows that the evolution of network activity can be separated into the evolution of *N* individual components,

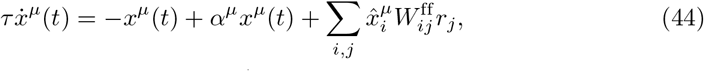

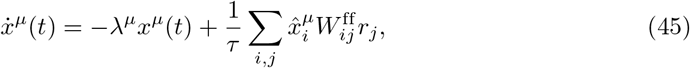

see, e.g., [41]. Here we defined

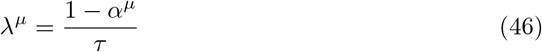

as the decay rate of network activity in the *μ*th eigenmode. The response time of the network equals 1*/λ*^*μ*^, if it is initialized in the *μ*th eigenmode. When initialized by *x*(0), the state decays to a stationary value in the different components,

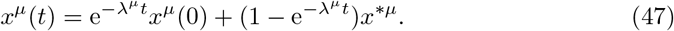

Here 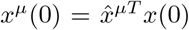 are the initial and 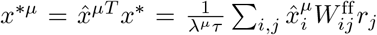 the final stationary values of the *μ*th component in the eigenbasis. The vector of the stationary dynamics *x*^∗^ has the entries 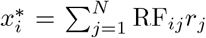, if the network generates the desired stationary dynamics. In the absence of recurrent input we have *W* ^rec^ = 0 and thus *α*^*μ*^ = 0, such that the activity decays at a rate of *λ* = 1*/τ* to its target. Positive eigenvalues *α*^*μ*^ *>* 0 mean a slower decay. At *α*^*μ*^ = 1 there is no decay at all, and for *α*^*μ*^ *>* 1 activity diverges. Stable activity thus requires the largest eigenvalue of the recurrent weight matrix to be smaller than one, such that together with the individual intrinsic decay of each neuron the dynamics are a contraction.

Because the system is linear, the different eigenmodes decay independently from each other at different rates, following eq. (47). Thus with time the faster-decaying modes connected to smaller eigenvalues become exponentially suppressed relative to the dominant, slowest-decaying mode.

The recurrent weight matrices in our recurrent networks eq. (5) furthermore have the property that they are circulant matrices, i.e. each row is equal to the row before rotated one element to the right. The eigenvalues of such matrices are given by the explicit formula [94]

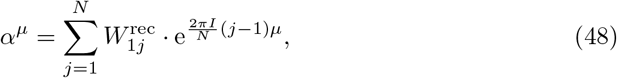

where *I* is the imaginary unit. For purely excitatory networks, we have 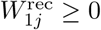, such that *α*^*μ*^ is maximal if *μ* = 0, as otherwise the real part of each addend in eq. (48) is smaller or equal. The corresponding eigenvector of *W* ^rec^ is

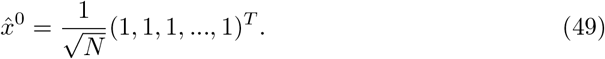

therefore, we obtain the slowest convergence for

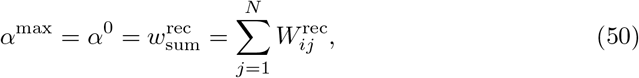

where 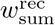 is the row sum of the recurrent coupling matrix, which is independent of *I* as the matrix is circulant. The slowest exponential decay dominates the behavior for longer times. Inserting *α*^max^ into eq. (46) thus yields *τ*_resp_, the generic time scale of convergence of the network dynamics eq. (5) to the target state 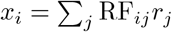,

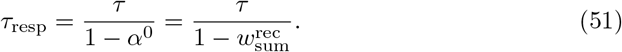

### S2 Loss evolution of excitatory networks

In this section, we compute the time evolution of the network loss for our networks without SFA and explicit inhibition. We assume that the networks receive constant input and have initial state *x*_*i*_(0) = 0. All convergence time scales that are present in the network (cf. eqs. (46) and (48)) could in principle contribute to this time evolution. We will however see, that the decay time of the loss is equal to that of the slowest-decaying mode.

We define the loss as the 1-norm of the deviation of the network activity from the target activity,

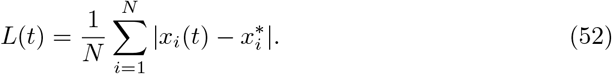

Under the assumption that network activity does not ‘overshoot’, i.e. that it is always lower than or equal to the target activity, 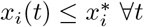, we can replace the loss function by a linear loss function

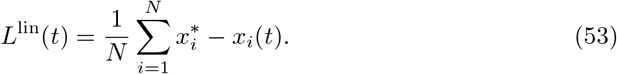

we can express the linearized loss as the (scaled) projection of the deviation of the activity from its target, *x*^∗^ − *x*(*t*), onto the eigenvector 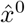 eq. (49),

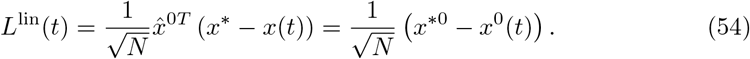

the linear loss thus has a single exponential decay. This is a consequence of the fact that (1, 1, 1, …1)^*T*^ is an eigenvector of the dynamics, which, in turn, holds because the row sum of a circulant matrix is the same for each row. The linear loss therefore decays with the time constant *τ*_resp_ obtained in eq. (51) to its stationary value, 0.

### S3 Cooperative coding minimizes the number of required synapses in 1D excitatory networks

The cooperatively coding recurrent networks eq. (5) and, with SFA, eq. (17) generate the required stationary dynamics eq. (1) with three synapses per neuron. Here we show that this is the minimal number of synapses. Specifically, we show that it is impossible to construct the RF of one neuron as the sum of only one or two other RFs and/or feedforward inputs.

Any network of the form eq. (4) that correctly implements the desired target response has RFs that satisfy eq. (8),

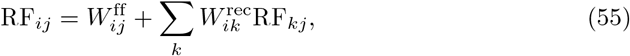

i.e. RF_*i*_ is a sum of localized peaks from feedforward inputs and extended two-sided exponentials from recurrent inputs (see Fig. 2c) for an example). First, there needs to be at least one (nonzero) feedforward synapse, as otherwise the network could not respond to input neuron activity. Clearly two feedforward synapses cannot solve the problem for RFs with *n*_RF_ *>* 2. But the sum of a localized peak from a feedforward input and the extended response from a recurrent input (with nonzero coefficients) cannot match the target shape (because the difference between the target RF and another RF is either zero or has extended support). Thus every neuron needs at least three synapses to implement the target network response.

### S4 Spike frequency adaptation

Here we study the dynamics of a 1D network with SFA, which are given by eq. (17). Fig. 8a) shows the dynamic response of neuron *j*_0_, whose preferred input is presented as isolated unit input. The initial dynamics are faster than for a network without SFA. This is because the adaptation current effectively reduces the membrane time constant and because it does not yet fully compensate for the stronger recurrent and feedforward weights, as it is slow (see eqs. (14) and (16)).

To quantify the change in response speed, we compute the integrated loss, as a fast convergence implies that the integrated loss is small. We normalize the loss, Loss(*t*) = |*x*(*t*) − *x*^∗^ |_1_*/* |*x*^∗^| _1_ to render it comparable for different RF widths. We define the integrated loss as the temporal mean of the normalized loss. Fig. 8d) shows the dependence of the integrated loss on the SFA time constant. The left-most data point at *τ*_SFA_ = 0 corresponds to the zero-lag limit where *u*_*i*_(*t*) = *x*_*i*_(*t*), which effectively modifies the leak current to −(1 + *a*_SFA_)*x*_*i*_(*t*) (eq. (16)), resulting in a reduced neuronal time constant *τ*→ *τ/*(1 + *a*_SFA_). Consequently, in this limit and for *a*_SFA_ = 1, the response time of the SFA network is half as large as that of a network without SFA, which is 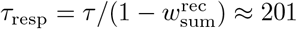, eq. (11). Also the integrated loss is smaller; without SFA it would be ≈ 0.413. The negative slope at *τ*_SFA_ = 0 and the clear minimum at *τ*_SFA_ ≈ 0.87 show that the network can increase its response speed beyond this effective reduction of *τ* by utilizing the larger recurrent and feedforward weights, eq. (15), which are not yet fully compensated during equilibration. For oscillating dynamics, *τ*_resp_ can jump discontinuously. This happens if the time point at which all later losses are below e^−1^ jumps from one oscillation peak to the next. Therefore we use the integrated loss here as the minimization target.

Fig. 8f) shows the integrated loss as a function of *a*_SFA_ and 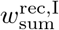 in networks that combine SFA and balancing inhibition. The integrated loss is minimized in networks with critical inhibitory strength and without SFA. Larger adaptation can partly compensate weaker inhibition; the data suggest that the optimal values of the adaptation and inhibition parameters satisfy the relation 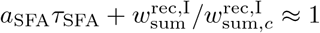.

### S5 Effective residual inhibitory interaction strength

The residual inhibitory interaction term in the dynamics eq. (22) reads

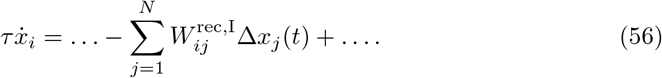

In the following we compute the effective strength of the interaction mediated by this term, which we define as the total, integrated contribution to the state change of feature neuron *i* that it causes. For this we assume that the presynaptic activity *x*_*j*_ of neuron *j* changes only for a limited amount of time, i.e. its derivative has limited support. Concretely we assume that 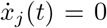 for |*t*| *> T* − *τ*_lag_ for some finite time *T*. The component of the overall state change *δx*_*i*_ = *x*_*i*_(*T*) −*x*_*i*_(−*T*) in neuron *i* between –*T* and *T* that is caused by the residual inhibitory term due to changes in neuron *j* is then given by

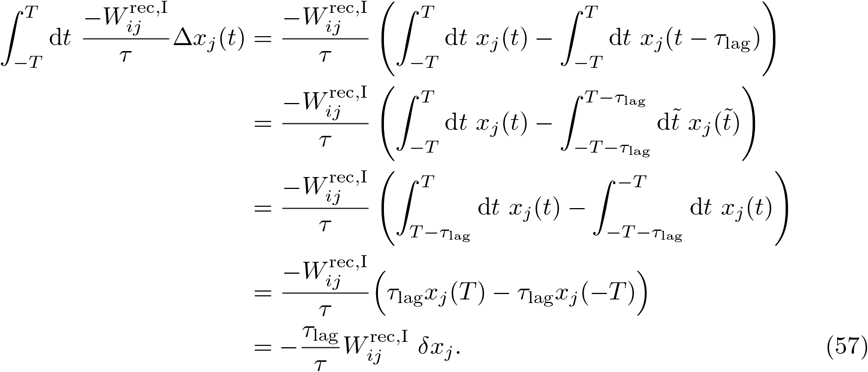

Here we substituted 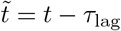 in the second line; in the third line we used that large parts of the two integrals cancel and in the fourth line that *x*_*j*_(*t*) is constant between *T* − *τ*_lag_ and *T* as well as between −*T* − *τ*_lag_ and −*T*. *δx*_*j*_ = *x*_*j*_(*T*) − *x*_*j*_(−*T*) is the total change in presynaptic activity. The result states that a change *δx*_*j*_ in presynaptic activity causes a total postsynaptic state change of 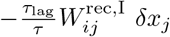. This may also be intuitively understood as follows: A step-like activity change in *x*_*j*_(*t*_0_) by *δx*_*j*_(*t*_0_) at time *t*_0_ increases ∆*x*_*j*_(*t*) by *δx*_*j*_(*t*_0_) for all *t* with *t*_0_ ≤ *t* ≤ *t*_0_ + *τ*_lag_. According to eq. (56) it thus changes 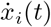 by 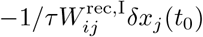 for a duration of *τ*_lag_. The integrated effect is thus 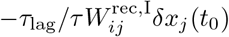. A continuous change of *x*_*j*_(*t*) may be seen as assembled of many small step-like ones, which together sum to *δx*_*j*_ and thus have the integrated effect eq. (57).

In the limit of small *τ*_lag_, the residual inhibitory interaction term transmits the temporal derivative of the neuronal dynamics in excitatory manner, adding a contribution with the same sign to the temporal derivative of the postsynaptic neuron. To see this, we scale the inhibitory weights with *τ*_lag_ in such a way that the total, integrated effect of the residual inhibition due to a presynaptic activity change at *t* is independent of the lag between excitatory and inhibitory activity, *τ*_lag_, i.e. we set 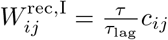 with constant *c*_*ij*_, compare eq. (57). In the limit of short lag the delayed interaction term then becomes 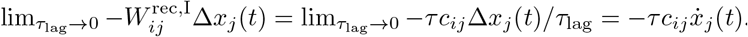. The prefactor −*τc*_*ij*_ *>* 0 is positive, which renders the coupling excitatory.

### S6 Loss evolution of balanced networks

To analytically estimate the evolution of the loss in our networks with balancing, delayed inhibition, we use again the linearized L1 loss eq. (53)), i.e. we assume again that the activity is always lower than or equal to the target activity. In the purely excitatory network, the linearized loss satisfied some simple dynamical equations, eq. (54) and eq. (45). We will see in the following that this also holds in our balanced networks. To obtain the dynamical equation we compute the temporal derivative of the loss, using the linear time evolution of the activityeqs. (22) and (24). We assume that the input is constant, *r*_*j*_(*t*) = *r*_*j*_, and use the knowledge that for such input the dynamics converge to a stationary target state 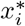. This yields

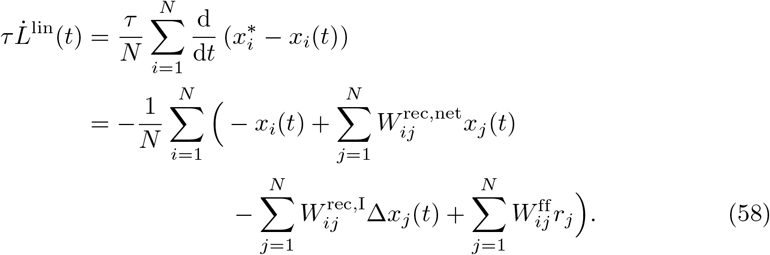

we now specialize the computation further by taking into account that in the cases of interest for us, *W* ^rec^, *W* ^rec,I^ and thus also *W* ^rec,net^, as well as *W* ^ff^ are circulant matrices. This implies that the column sums 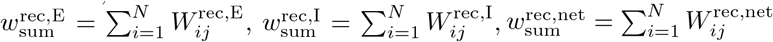 and 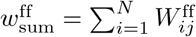 are independent of the column *j*. Equation (58) thus simplifies to

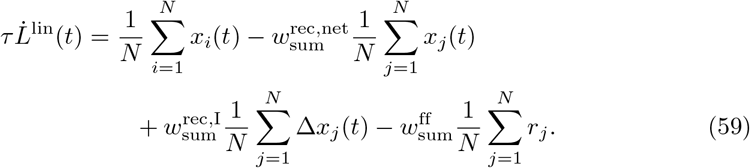

the networks that we want to track analytically start with *x*_*i*_(0) = 0, such that the initial linear loss is 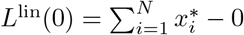. To describe the network loss dynamics, we can thus use

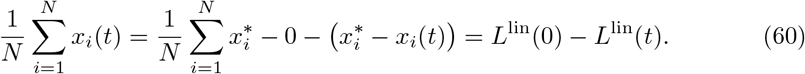

An alike equation holds for 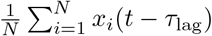, such that

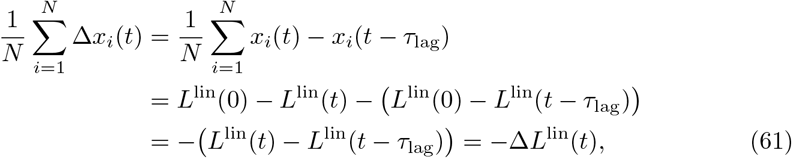

where we introduced the abbreviation

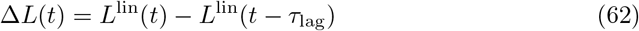

for the difference between the current and the delayed loss. Inserting eq. (60) and eq. (61) into eq. (59) gives

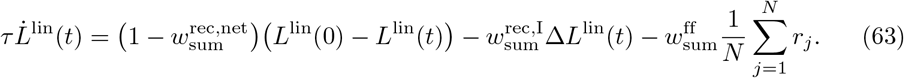

to eliminate the explicit occurrence of the inputs, we use that in the stationary state, which is reached for *t* → ∞, we have *L*^lin^(∞) = 0, 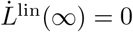 and ∆*L*^lin^(∞) = 0. For *t* → ∞, eq. (63) thus shows that

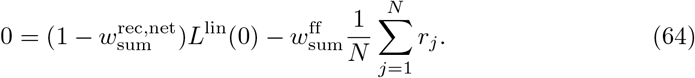

Employing this in eq. (63) gives our final dynamical equation for the linearized loss in terms of the variable *L*^lin^ only,

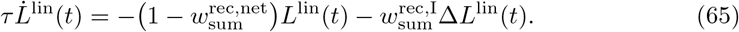

we solve this delay differential equation with the ansatz

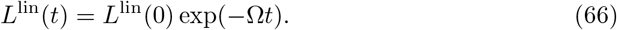

we note that

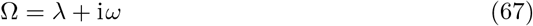

is generally complex. The ansatz implies

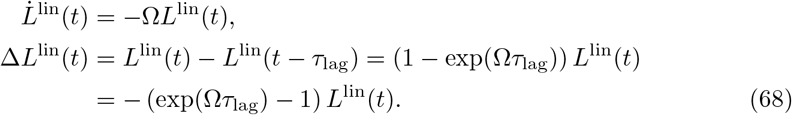

Inserting eq. (68) into eq. (65) and dividing by −*τ L*^lin^(*t*) yields

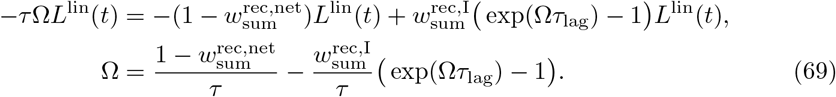

we see immediately that in the absence of inhibition, 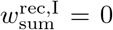, we have 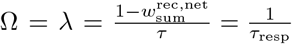 (cf. eq. (11)), such that the decay rate of the purely excitatory network eq. (4) is recovered, as it has to be. To solve eq. (69) in presence of inhibition, we rewrite it as

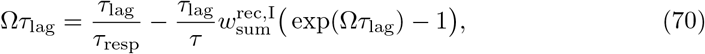

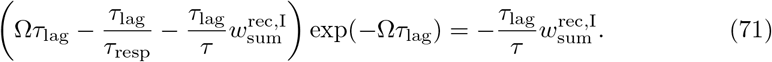

Substituting

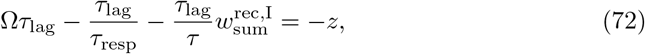

we obtain

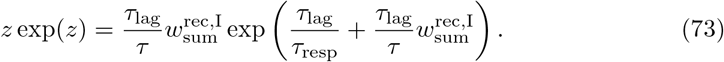

the branches of the Lambert W function solve *z* exp(*z*) = RHS for *z*. Applying them to eq. (73) yields *z* = *W*_*k*_(RHS), where *W*_*k*_ denotes the *k*th branch and RHS the right hand side of eq. (73),

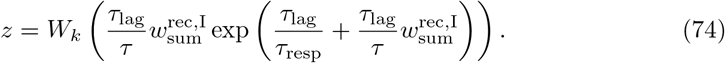

Resubstituting for *z* then gives the decay rate *λ* (cf. eq. (67)) and, if an oscillation is present, the oscillation frequency *ω* of the linearized loss,

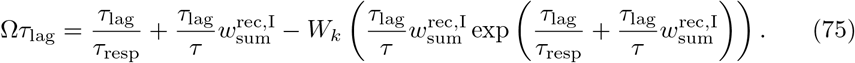

Figs. 5 and 9 show the relevant solutions, obtained from the branches *k* = 0 and *k* = − 1.

Similarly to the dynamics of a dampened harmonic oscillator, there is a transition from an exponentially-decaying (‘overdamped’) regime with two decay rates to an oscillating (‘underdamped’) regime with a single one. At the transition point we have critical inhibitory strength (‘critical damping’). In the oscillatory regime it is the amplitude of the oscillation that decays exponentially. Because the actual error is always smaller or equal to the amplitude, having slightly larger than critical inhibition may be optimal, see Fig. 5a). The solutions corresponding to branches other than *k* = 0, − 1 have markedly higher decay rates and are thus of little relevance. They also have high oscillation frequencies with periods smaller than the lag.

We note that in the case of oscillations (*ω* ≠ 0), the linearized loss eq. (66) periodically reaches zero. This corresponds to 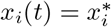 for all neurons, meaning that all neurons simultaneously go from positive to negative deviations from their target activities, or vice-versa. In the full network model, however, these transitions will occur at different times, so that when one neuron matches its target activity there are others that don’t. Therefore, the minima of the oscillation are not at zero but at a final error, see Fig. 5a).

We now determine the critical inhibitory strength and the critical decay rate. These are determined by *z* = *W*_*k*_(RHS) (eq. (74), with RHS the right hand side of eq. (73)) having only one real solution *z*. This happens at the branch point RHS = −e^−1^ where the 0th and −1st branch agree, *z* = *W*_0_(−e^−1^) = *W*_−1_(−e^−1^) = −1 [95]. We first set RHS = *z*e^*z*^ = −e^−1^ to obtain 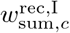,

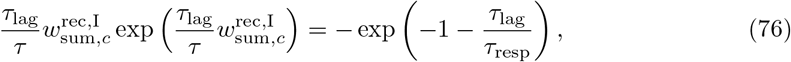

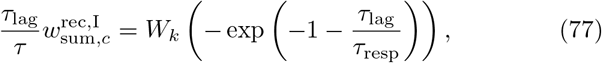

where we have again used a (yet unspecified) branch of the Lambert W function to solve for 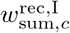. We know that the inhibitory strength must be real and negative; this can only hold for *k* = 0 or *k* = − 1. It might seem surprising that we obtain two solutions here. The reason is that there are actually two critical points, seeFig. 9a): one with a positive decay rate, and another with a negative decay rate, which thus describes exponential growth. We are interested only in the converging dynamics and thus choose *k* = 0.

Next we resubstitute 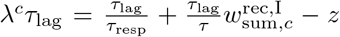 (eq. (72) with Ω replaced by the critical, real decay rate *λ*^*c*^), use *z* = − 1 and insert the critical inhibitory strength eq. (77) to obtain the critical decay rate,

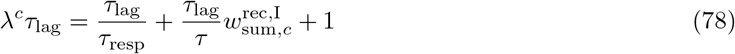

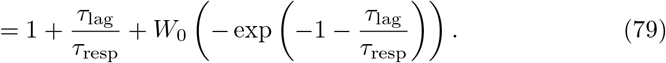

the response time of the balanced network is the inverse of *λ*^*c*^,

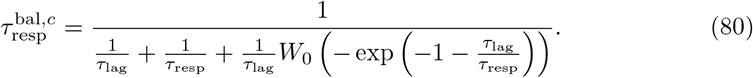

we now want to derive a more easily interpretable approximation for *λ*^*c*^ and 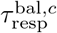 for small *τ*_lag_*/τ*_resp_. To this end, we modify eq. (79) to

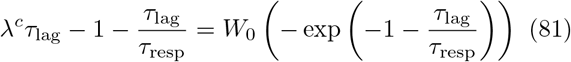

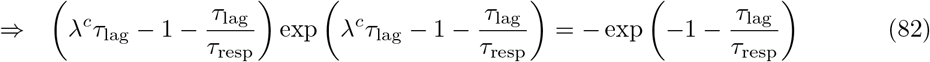

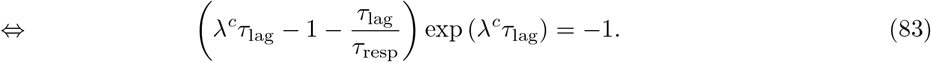

Here we have applied the inverse of *W*_0_, *z* → *z*e^*z*^, to both sides, and multiplied with exp(−1 −*τ*_lag_*/τ*_resp_). Next we assume that *λ*^*c*^*τ*_lag_ is small, meaning that the response time is much larger than the E-I lag, expand the exponential up to the second power in *λ*^*c*^ and *τ*_lag_, and solve for *λ*^*c*^*τ*_lag_,

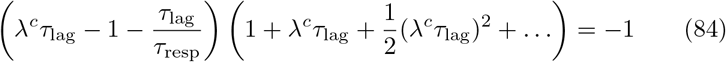

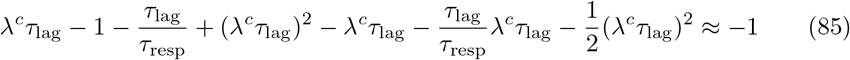

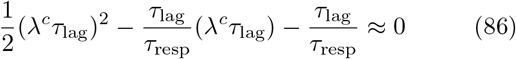

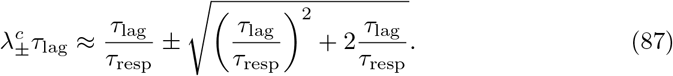

Here only the positive solution makes sense. Since the delayed inhibition speeds up responses, our assumption that *λ*^*c*^*τ*_lag_ is small implies that also *τ*_lag_*/τ*_resp_ with the response time of the network without delayed inhibition is small. We can thus neglect in the radicand of eq. (87)’s RHS the quadratic term compared to the linear one. Compared to the resulting square root term we can neglect the first, linear RHS term, such that we obtain

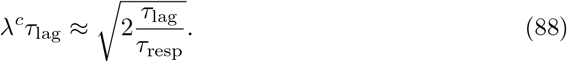

the response time of the balanced network follows as the inverse of *λ*^*c*^,

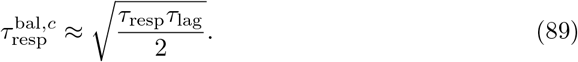

Fig. 9 shows that, despite the simple formula, the quality of the approximation is very good.

### S7 Initial response of balanced networks

In the following we explain the discrepancy between the analytical and numerical linearized loss evolution, which results in different response times (see Fig. 6a), c), stems from different initializations of neuronal activity. For this, we first note that an exponential function eq. (66), *L*^lin^(*t*) = *L*^lin^(0) exp(−Ω*t*), with Ω solving eq. (69) and arbitrary amplitude *L*^lin^(0), is an eigenmode of the time evolution of the linearized loss eq. (65), as the time evolution preserves its functional form. The results of the last section describe the loss evolution if the system is in such an eigenmode. However, in the main text simulations, networks are not initialized in an eigenmode: They are initialized with zero activity and respond to a sudden jump in input. This corresponds to an initial state with *x*(*t*) = 0 for *t* ≤ 0; in particular ∆*x*(0) = 0. Therefore both the initial and earlier losses equal the maximal loss, *L*(*t*) = *L*(−*τ*_lag_) = |*x*^∗^ |_1_*/N* for *t* ≤ 0 and ∆*L*(0) = 0, see eqs. (61) and (62) and Fig. 10b). In contrast, for the eigenmodes we have ∆*L*^lin^(*t*) = − (exp(Ω*τ*_lag_) − 1)*L*^lin^(*t*) for *t*≤ 0 (from eqs. (62) and (66)); especially ∆*L*^lin^(0) *<* 0. The loss is thus not initialized in an eigenmode; but the decay rate of the loss converges to that of the slowliest decaying eigenmode over time.

**Fig 10:**
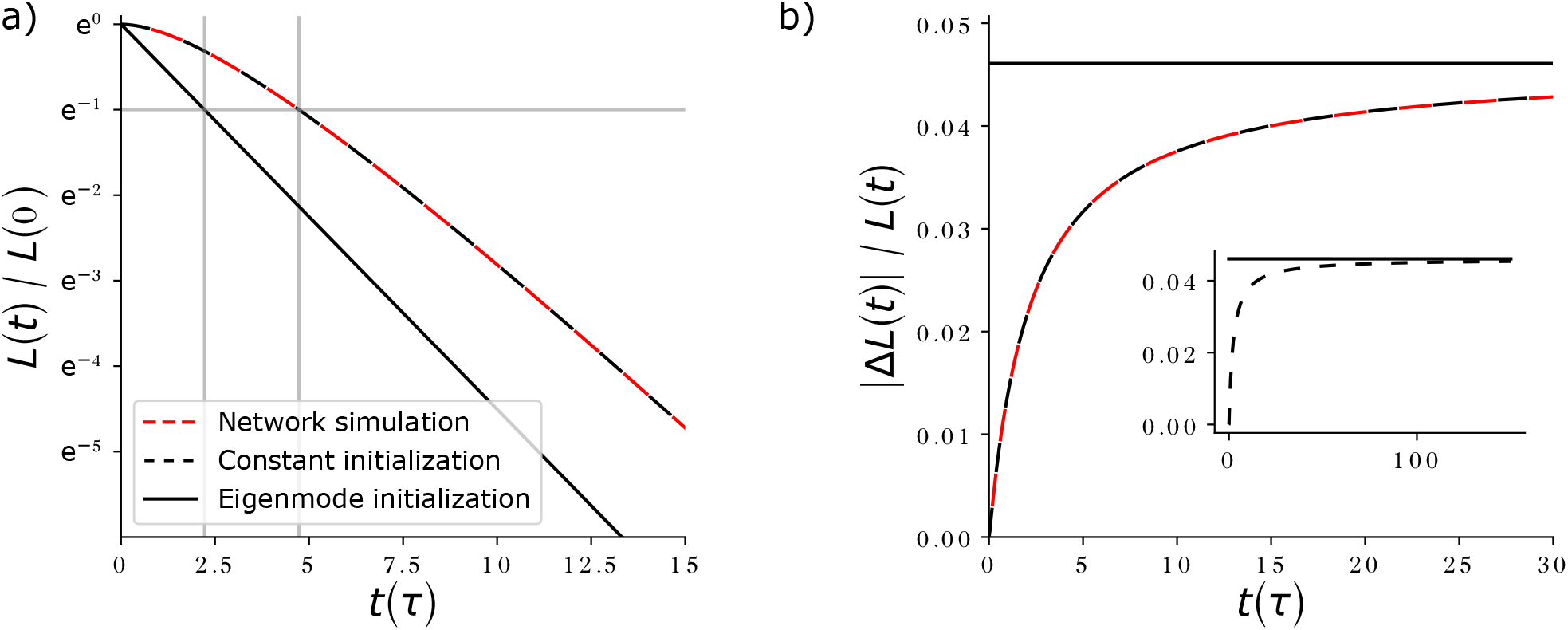
Effects of initial conditions on loss evolution. **a)** Normalized loss evolution obtained by solving eq. (65) (black curves) for critical inhibitory strength. The loss is either initialized in the slowest eigenmode (solid, analytical solution eqs. (66) and (69)), like in our analytical computations, or it is initialized as a constant function (dashed, *L*(*t*) = *L*(0) for *t* ≤ 0, numerical solution), like in our network simulations. Vertical lines indicate the time at which the error drops below e^−1^, which experimentally defines the response time. The theory with constant initialization describes the loss evolution of a numerically simulated network (red dashed, *N* = 200) well. With time, the loss decay rate approaches the same value, independent of initialization; the curves become parallel. **b)** In the for the chosen parameters non-oscillatory slowliest decaying eigenmode, ∆*L*(*t*) is always proportional to *L*(*t*) (solid horizontal line at the proportionality factor 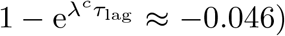. With constant initialization (dashed curves), ∆*L*(*t*) is initially zero, such that the impact of the interactions mediated by *W* ^rec,I^ (see eq. (24)) is initially small. ∆*L*(*t*) then tends to the same proportionality to *L*(*t*) as for the eigenmode initialization. Inset: loss evolution for long times. Parameters: 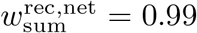 *τ* = 1, *τ*_lag_ = 0.1, 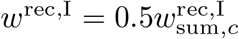; we thus have 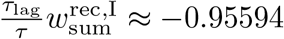; the displayed slowliest decaying eigenmode is non-oscillatory, cf. Fig. 9.

For the chosen initial conditions the loss decay speeds up during this process, see Fig. 6c and Fig. 10a),b). This can be understood as follows: In our network simulations, activity initially increases, such that ∆*x*_*i*_(*t*) is positive. The amplifying current term 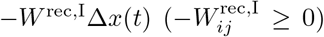 in eq. (58) is thus positive, which causes a dynamical speed-up in the activity increase.. Its sum is proportional to −∆*L*(*t*), which is initially zero. The term 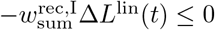 introduces the dynamical speed up evoked by ∆*x*_*i*_(*t*) into the equation of the loss dynamics, where it causes a faster decay of the loss. As both ∆*x*_*i*_(*t*) and |∆*L*(*t*) | increase, the terms that generate the speed up in the activity and in the loss dynamics increase in absolute value. As a consequence, the activity increases faster and the loss decays faster. The inset in Fig. 10b) shows that |∆*L*| increases (initially in absolute, later in relative terms) and approaches the same proportionality to *L* as for the slowliest decaying eigenmode, which is non-oscillatory in the displayed example. This means that the system settles in the eigenmode. The loss curves then become parallel lines on the logarithmic axes (extrapolation of Fig. 10a)).

This different initialization and the resulting initially lower impact of lagged interactions explain why the red data points in Fig. 6 show longer response times than expected from the theory.

